# Normative high-frequency oscillation phase-amplitude coupling and effective connectivity under sevoflurane

**DOI:** 10.1101/2025.03.18.644050

**Authors:** Ethan Firestone, Hiroshi Uda, Naoto Kuroda, Kazuki Sakakura, Masaki Sonoda, Riyo Ueda, Yu Kitazawa, Min-Hee Lee, Jeong-Won Jeong, Aimee F. Luat, Michael J. Cools, Sandeep Sood, Eishi Asano

## Abstract

Resective surgery for pediatric drug-resistant focal epilepsy often requires extraoperative intracranial electroencephalography recording to accurately localize the epileptogenic zone. This procedure entails multiple neurosurgeries, intracranial electrode implantation and explantation, and days of invasive inpatient evaluation. There is a need for methods to reduce diagnostic burden and introduce objective epilepsy biomarkers. Our preliminary studies aimed to address these issues by using sevoflurane anesthesia to rapidly and reversibly activate intraoperative phase-amplitude coupling between delta and high-frequency activities, as well as high-frequency activity-based effective connectivity. Phase-amplitude coupling can serve as a proxy for spike-and-wave discharges, and effective connectivity describes the spatiotemporal dynamics of neural information flow among regions. Notably, sevoflurane activated these interictal electrocorticography biomarkers most robustly in areas whose resection led to seizure freedom. However, they were also increased in normative brain regions that did not require removal for seizure control. Before using these electrocorticography biomarkers prospectively to guide resection, we should understand their endogenous distribution and propagation pathways, at different anesthetic stages.

In the current study, we highlighted the normative distribution of delta and high-frequency activity phase-amplitude coupling and effective connectivity under sevoflurane. Normative data was derived from nineteen patients, whose ages ranged from four to eighteen years and included eleven males. All achieved seizure control following focal resection. Electrocorticography was recorded at an isoflurane baseline, during stepwise increases in sevoflurane concentration, and also during extraoperative slow-wave sleep without anesthesia. Normative electrode sites were then mapped onto a standard cortical surface for anatomical visualization. Dynamic tractography traced white matter pathways that connected sites with significantly augmented biomarkers. Finally, we analyzed all sites —regardless of normal or abnormal status — to determine whether sevoflurane-enhanced biomarker values could intraoperatively localize the epileptogenic sites. We found that normative electrocorticography biomarkers increased as a function of sevoflurane concentration, especially in bilateral frontal and parietal lobe regions (Bonferroni-corrected p-values <0.05). Callosal fibers directly connected homotopic Rolandic regions exhibiting elevated phase-amplitude coupling. The superior longitudinal fasciculus linked frontal and parietal association cortices showing augmented effective connectivity. Higher biomarker values, particularly at three to four volume percent sevoflurane, characterized epileptogenicity and seizure-onset zone status (Bonferroni-corrected p-values <0.05). Supplementary analysis showed that epileptogenic sites exhibited less augmentation in delta-based effective connectivity. This study helps clarify the normative distribution of, and plausible propagation pathways supporting, sevoflurane enhanced electrocorticographic biomarkers. Future work should confirm that sevoflurane-activated electrocorticography biomarkers can predict postoperative seizure outcomes in larger cohorts, to establish their clinical utility.

## Introduction

Millions of children are impacted by focal epilepsy, and prompt treatment is needed to avoid cognitive comorbidities.^1–3^ Around 30 percent of these patients will develop drug-resistant epilepsy that often requires surgery to remove the epileptogenic zone responsible for generating habitual seizures.^3–6^ Since localizing the epileptogenic zone using noninvasive modalities, alone, can be challenging, many patients require intracranial EEG (iEEG) electrode implantation and days of extraoperative iEEG monitoring to capture spontaneous ictal events and identify the origin of these discharges. ^7–15^ The two-stage procedure involves significant risk factors from multiple neurosurgeries and chronic brain implants.^11,16–17^ It is thus ideal to acutely record intraoperative electrocorticography (ECoG) directly from the cortical surface, immediately followed by focal resection.^18–19^ Moreover, conventional methods rely on visual inspection of iEEG traces to identify epileptiform signals, such as spike-and-wave discharges. This is subject to inter-rater variability and makes it difficult to compare relative levels of epileptogenicity across sites.^20–21^ Hence, better intraoperative techniques using inducible, objective ECoG measures are needed to avoid these pitfalls.

Sevoflurane may offer a unique solution because it rapidly and reversibly augments interictal epileptiform activity, unlike other general anesthetics which suppress such signals.^22–39^ Furthermore, recent studies have shown that sevoflurane enhances objective ECoG epilepsy biomarkers, including the occurrence rates of high-frequency oscillations (HFOs) at ≥80 Hz, phase-amplitude coupling between the 3-4 Hz delta phase and HFO amplitude (delta–HFO phase-amplitude coupling: delta-HFO PAC), and HFO-based effective connectivity (EC), particularly in epileptogenic regions.^40–72^ Delta–HFO phase-amplitude coupling is considered a reliable proxy for interictal spike-and-wave discharges.^57–59,62^ Thus, we expected that sites with elevated values might serve as key nodes within networks where epileptiform signals originate and propagate. We also expected that HFO-based effective connectivity might characterize the spatiotemporal dynamics of high-frequency activity propagation within such networks.^69–71^ However, our recent investigations demonstrated that sevoflurane, likewise, increased delta-HFO phase-amplitude coupling and HFO-based effective connectivity in normative brain regions that need not be resected for seizure freedom.^68–69^ These results highlight the need to determine the endogenous distribution of ECoG biomarkers across different sevoflurane levels. Additionally, identifying the structural pathways that mediate sevoflurane-induced signal propagation is valuable. We believe that characterizing the expected response in the normal brain will facilitate the use of sevoflurane anesthesia for intraoperative localization of the epileptogenic zone.

As such, we analyzed a limited pediatric ECoG dataset to begin deducing the normative distribution of interictal delta-HFO phase-amplitude coupling (quantified by modulation index^62–66^) and HFO-based effective connectivity (quantified by transfer entropy^70–71^), at various anesthetic stages. We next employed dynamic tractography to trace major white matter pathways that directly connected cortical sites showing significant ECoG biomarker augmentation. Finally, we determined the fidelity of using our objective, intraoperative ECoG biomarkers for characterizing epileptogenicity and seizure onset zone (SOZ) status under sevoflurane. We hypothesized that sevoflurane would significantly augment delta-HFO phase amplitude coupling and HFO effective connectivity, in normative electrode sites. We further postulated that cortical sites showing the greatest degree of co-augmentation would be linked by underlying white matter fibers. We finally hypothesized that sevoflurane would elevate these objective, intraoperative ECoG biomarkers most in epileptogenic and SOZ sites.

## Materials and methods

### General methods

We investigated intraoperative, interictal ECoG data from 19 pediatric, drug-resistant focal epilepsy patients that underwent resective surgery at the Children’s Hospital of Michigan (Detroit, MI, USA) and achieved ILAE class-I seizure outcomes. This study was strictly observational; all data was collected during standard-of-care treatment for drug-resistant focal epilepsy, with no deviation for scientific curiosity. The results were not available to influence surgical plans. Two-stage patients *(n = 11)* went through the following clinical course: [1] non-invasive presurgical evaluation,^9^ [2] implantation of intracranial electrodes in conjunction with sevoflurane-activated, intraoperative ECoG, [3] 3-5 days of extraoperative functional brain mapping via iEEG recording, [4] a second neurosurgery for explantation of intracranial electrodes immediately followed by focal cortical resection, and [5] one-year post-operative follow-up to determine seizure outcome status. The study also included single-stage *(n = 8)* patients who received: [1] non-invasive presurgical evaluation, [2] acute, intraoperative ECoG recording, immediately followed by focal cortical resection, and [3] one-year postoperative follow up to determine seizure status. The statistical models addressed potential bias between one-versus two-stage surgery. Importantly, all 19 patients experienced intraoperative ECoG recording in the presence of stepwise-rising sevoflurane. This procedure is standard care at Children’s Hospital of Michigan to optimize intracranial electrode sampling of implicated brain regions for invasive, extraoperative iEEG recording.

We retrospectively employed signal processing on the intraoperative ECoG and extraoperative iEEG data. We calculated delta-HFO PAC (rated by modulation index) and HFO EC (rated by transfer entropy) under stepwise increasing sevoflurane (2-4 vol%), isoflurane (1 vol%), and during slow wave sleep; the latter two conditions were redundant controls. A linear mixed model tested whether normative delta-HFO PAC and/or HFO EC increased as a function of sevoflurane concentration. Normative electrode sites were defined as those outside the resection area, the SOZ,^9^ areas generating interictal epileptiform discharges,^61^ and regions with MRI-visible lesions.^73,74^ Pooled, normative electrodes were then interpolated onto a standard cortical template to visualize the anatomical distribution of delta-HFO PAC and HFO EC, at each anesthetic stage. Diffusion-weighted imaging (DWI) delineated white matter streamlines connecting site-pairs with co-augmented ECoG biomarkers, in “dynamic tractography”.^66^ Next, we included all electrode sites, regardless of normality, to deduce if relatively greater sevoflurane-induced augmentation of delta-HFO PAC and/or HFO EC could determine the epileptogenic status of electrodes. The epileptogenic zone was defined as the resected cortical area, in patients who achieved one-year postoperative seizure freedom.^69^ Supplementary analysis tested if the biomarkers could likewise characterize the SOZ^9^: the area of ictogensis within the epileptogenic zone that is prospectively defined by board-certified clinical neurophysiologists. Since spike-and-wave discharges characteristically contain a 3-4 Hz component and delta waves are the other variable in PAC,^57,59,61,62^ we also created a supplementary normative distribution, along with predictions models, for 3-4 Hz delta EC (rated by transfer entropy).

A final supplementary analysis tested whether more complete resection of high-value ECoG biomarker sites could predict ILAE seizure outcomes. This analysis included an extra four patients (aged 6-20 years; 3 males) who underwent two-stage surgery, including sevoflurane-based ECoG and extraoperative iEEG recordings, but failed to achieve ILAE class-I seizure outcome. Their detailed clinical information can be found in Supplementary Table 1.

### Patient population

The study included 19 pediatric patients (aged 4-18 years; 11 males) who received resective epilepsy surgery and achieved one-year post-operative ILAE-defined seizure freedom. We obtained written consent from all patients or legal guardians of those under 18 years old and from those who were unable to provide their own consent. Detailed clinical characteristics can be found in Table 1. All analyses were approved by the Institutional Review Board (IRB) at Wayne State University, Detroit MI, USA.

#### Inclusion Criteria

[1] children, aged 4-20 years, with a diagnosis of drug-resistant focal epilepsy; [2] underwent resective epilepsy surgery; [3] intraoperative ECoG was recorded under isoflurane maintenance and a short period of sevoflurane.

#### Exclusion Criteria

[1] malignant brain tumor; [2] progressive neurodegenerative or metabolic disorder; [3] presence of a major developmental brain malformation which can confound anatomical landmark identification; [4] previous epilepsy surgery.

### Implantation of intracranial electrodes and intraoperative ECoG

Intracranial electrodes were implanted per standard clinical management of drug-resistant focal seizures.^9,62^ First, a multidisciplinary group of board-certified clinicians synthesized non-invasive neuroimaging, neurophysiology, and neuropsychological data to estimate the location of the epileptogenic zone. Pediatric neurosurgeons then implanted intracranial electrodes on the implicated brain hemisphere(s) to be subsequently used for extraoperative, iEEG functional brain mapping. Anesthesia was induced with propofol, fentanyl, and rocuronium. Additional doses of fentanyl and rocuronium were administered as needed, and thereafter, 60% oxygen or above was given throughout the surgery. During the craniotomy and intracranial electrode implantation, anesthesia was maintained with isoflurane (1 vol%). Patients were implanted with either surface platinum disk electrodes (10 mm center-to-center), depth electrodes (5 mm center-to-center), or a combination of the two. The number and spatial extent of iEEG electrodes were strictly based on clinical needs, with no deviation for scientific curiosity. After electrode placement, intraoperative ECoG recording was initiated, and the end-tidal carbon dioxide level was maintained below 40 mmHg. ECoG signals were recorded using a 1,000 Hz sampling rate and a bandpass filter at 0.016 – 300 Hz. Anesthesia was then switched to sevoflurane for a total of 15 minutes to temporarily and reversibly augment interictal epileptiform discharges, thereby reducing the risk of sampling errors. The sevoflurane concentration was progressively increased from 2 vol% to 3 vol%, and finally to 4 vol%, in a stepwise manner. These doses are clinically acceptable.^28,32,68,75^ The clinical neurophysiologist and anesthesiologist constantly communicated during the intraoperative ECoG recording. When a reduction in blood pressure was encountered and anticipated to be significant, the anesthesiologist mitigated sevoflurane’s effect on blood pressure by administering fluid boluses and vasopressors such as phenylephrine or epinephrine and proactively ceased increasing the sevoflurane level, as needed, after communication with the clinical neurophysiologist. A board-certified clinical neurophysiologist interpreted the intraoperative ECoG traces, in real-time, and communicated findings to the neurosurgeon who placed additional iEEG electrodes as needed. Anesthesia was then switched back to isoflurane (1 vol%) maintenance for the remainder of the surgery.

### Extraoperative iEEG and focal cortical resection

After implantation of intracranial iEEG electrodes, patients were transferred to the epilepsy monitoring unit for 3-5 days of extraoperative iEEG recording to delineate the epileptogenic zone and eloquent cortex defined by electrical stimulation mapping.^9,62,76^ Intracranial EEG signals were recorded using a 1,000 Hz sampling rate and a bandpass filter at 0.016 – 300 Hz. Each patient’s electrodes were co-registered to their reconstructed MRI, as previously reported.^62,77–78^ Board-certified clinical neurophysiologists used this data to determine the location of the SOZ.^9^ Focal resection was then carried out by a neurosurgical team aiming to remove the presumed epileptogenic zone (i.e., SOZ and neighboring MRI lesions), while preserving eloquent cortex to avoid creating sensorimotor and/or cognitive deficits. In the single-stage patients, acute, intraoperative ECoG was recorded during the same step-wise sevoflurane paradigm described above, to confirm resection margins. This was followed immediately by focal resection.

### Definition of anesthetic stages

Intraoperative ECoG recordings were split into the following 5-minute anesthetic stages for analysis: [1] isoflurane 1 vol% (Iso), [2] sevoflurane 2 vol% (Sev2), [3] sevoflurane 3 vol% (Sev3), [4] sevoflurane 4 vol% (Sev4). The eleven two-stage patients also included [5] extraoperative iEEG without anesthesia during slow wave sleep (SWS). Each anesthetic condition was further subdivided into one-minute epochs (e.g., ‘Sev2_1’ refers to the first minute of sevoflurane 2 vol%). All anesthetic concentrations were within clinical limits.^75^ Conditions [1] and [5] were redundant controls.

### Definition of normative, epileptogenic, and seizure onset zones

We analyzed intracranial EEG data from a total of *n = 1,608* electrode sites (isoflurane control) and *n = 1,344* (slow wave sleep control). Note, both controls involved the same electrode pool. The different electrode counts in isoflurane versus slow wave sleep reflected the different number of patients between combined one- and two-stage surgery patients (i.e., isoflurane control) versus only two-stage surgery patients (i.e., slow wave sleep control). These separate analyses were conducted to utilize redundant controls, because only the two-stage surgery patients underwent extraoperative iEEG recording without anesthesia, during slow wave sleep. Since all patients achieved ILAE class-I seizure freedom, electrode sites that were retained in the brain, outside of the SOZ and MRI-lesions, and free from interictal epileptiform discharges were retrospectively considered ‘normative’, as done previously.^66,73,74^ This included *n = 963* normative sites (isoflurane control) and *n = 816* normative sites (slow wave sleep control). For the prediction models, resected electrode sites in patients who achieved class-I seizure freedom were deemed ‘epileptogenic’ (isoflurane control: *n = 475* | slow wave sleep control: *n = 441*), while those retained in the brain were ‘non-epileptogenic’ (isoflurane control: *n = 1,133* | slow wave sleep control: *n = 903*), as done previously.^68,69^ Intraoperative photographs confirmed electrode classifications, as done previously.^62^ In two-stage patients, a board-certified, clinical neurophysiologist prospectively identified areas initially exhibiting sustained rhythmic iEEG discharges, accompanied by subsequent clinical seizure activity, which were not explained by state changes; such brain areas were assigned as ‘seizure onset zone’ (SOZ: *n = 76*).^72^ The remaining electrode sites were classfied as ‘non-seizure onset zone’ (non-SOZ: *n =1,268*).

### Electrophysiology and imaging analysis workflow

This study combined physiologic and anatomical workflows, allowing for precise localization of ECoG/iEEG signals (Fig. 1). The physiological arm began with raw ECoG/iEEG recordings under various anesthetic stages. Signals were preprocessed and time-frequency transformed to yield the spectral amplitude (square root of power) of HFO (80-300 Hz) and delta (3-4 Hz) frequency bands. These measures were then fed into modulation index^62,68,79^ and transfer entropy algorithms,^69–71^ calculating phase-amplitude coupling and effective connectivity, respectively. On the anatomical arm, electrode positions were interpolated onto 3-dimensional MRI scans of each individual patient’s brain. Electrodes were then spatially normalized and pooled onto standard cortical surface and white matter templates. Joining the two workflows in dynamic tractography^66,69^ created spatially accurate cortical maps of ECoG/iEEG biomarker activity, along with white matter tracts connecting electrophysiological hotspots.

**Figure 1.**
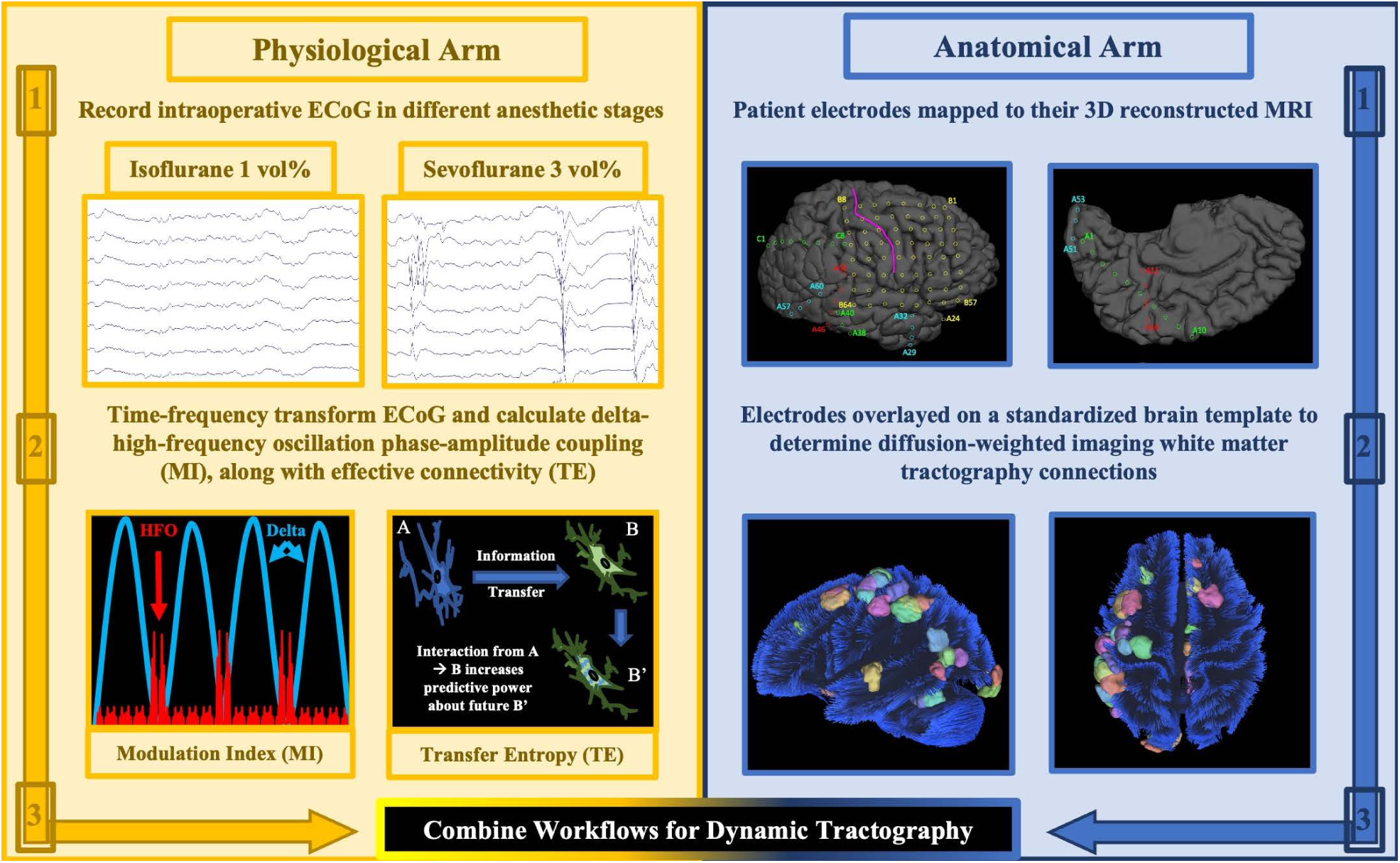
Electrophysiology and imaging analysis workflow. Physiologic (*left*) and anatomic (*right*) arms are combined to determine the group-level distribution of normative delta (3-4 Hz) and high-frequency oscillation (HFO: 80-300 Hz) phase-amplitude coupling (quantified by modulation index: MI) and effective connectivity (quantified by transfer entropy: TE). Dynamic tractography delineates white matter streamlines connecting electrophysiologic cortical hoptspots. Intraoperative ECoG = intraoperative electrocorticography.

### ECoG/iEEG preprocessing and time-frequency transformation

Investigators blind to the epileptogenic status of electrodes bandpass filtered the raw ECoG/iEEG from 0.016 – 300 Hz, notch-filtered 60 Hz noise, and exported the traces onto a bipolar montage. Preprocessing and time-frequency transformation were conducted using FieldTrip (https://www.fieldtriptoolbox.org/)^80^ open-source software, via Matlab R2022b (MathWorks, Natick, MA, USA). Visual screening removed channels and recording epochs polluted with artifacts. The wavelet method calculated power spectra of HFO (80-300 Hz) and delta waves (3-4 Hz). Taking the square root of power produced a spectral amplitude time-series.^69^ The spectral amplitude of each individual electrode and time-point was z-scored based on the given channel’s mean and standard deviation during the isoflurane control period. In two-stage patients, a separate, redundant dataset was made by z-scoring via the slow wave sleep control.

### Calculating phase-amplitude coupling via modulation index

Modulation index was calculated, as done previously.^62,66,68,79^ The cleaned, raw bipolar ECoG/iEEG data was fed into the open-source winPACT toolbox of EEGLAB (https://sccn.ucsd.edu/wiki/WinPACT) to compute modulation index.^79,81^ This software tool Hilbert transforms an EEG time series to quantify the strength of coupling between the instantaneous phase of delta waves (3-4 Hz) and HFO (80-300 Hz) amplitude. For each channel, PAC was calculated locally to determine the degree of coupling between 3-4 Hz delta phase and HFO amplitude, at that specific site. We chose to investigate HFO coupling exclusively with 3-4 Hz delta waves because previous studies from independent groups demonstrated that this specific narrow delta band is most representative of interictal spike- and-wave discharges.^57–58,66^

### Calculating effective connectivity via transfer entropy

Transfer entropy was calculated, as done previously.^69–71^ The z-scored spectral amplitude time series data was first time-binned (each bin at least 6 wave-cycles long). For a given bin, if the spectral amplitude rose above a z-score of 2 for at least 3 consecutive wave-cycles, then it was considered a ‘1’; otherwise, it was considered a ‘0’. The binary time series was fed through the transfer entropy algorithm to quantify effective connectivity between sites.^69–71^ All connections (efferent-emanating and afferent-receiving) for a given channel were averaged to get a single value for analysis. Transfer entropy-based effective connectivity was computed as a global measure, considering connections to all other channels, for a given site. Supplementary analysis probed possible differences between efferent and afferent.

### Anatomical localization of electrode sites

Anatomical localization of electrode sites was performed, as previously done.^62,69,77–78^ Board certified neurosurgeons used peri-operative photographs to interpolate each patient’s electrodes onto their 3D reconstructed MRI brain image. FreeSurfer software (https://surfer.nmr.mgh.harvard.edu/) spatially normalized electrodes onto the FreeSurfer average cortical template.^66,82–83^ This allowed group-level anatomical visualization of ECoG/iEEG biomarkers. Although the mathematical mixed models utilized data from all 19 patients (i.e., one- and two-stage patients), the cortical biomarker maps were derived from the eight two-stage patients that received ECoG recording under every anesthetic stage, as well as iEEG during slow wave sleep. Figure 2 shows the anatomical distribution of subdural electrodes from the subset of eight patients (*n = 431* normative electrodes). Biomarker values from the three, individual one-minute recording segments, at a given anesthetic stage, were averaged to produce one cortical map per condition.

**Figure 2.**
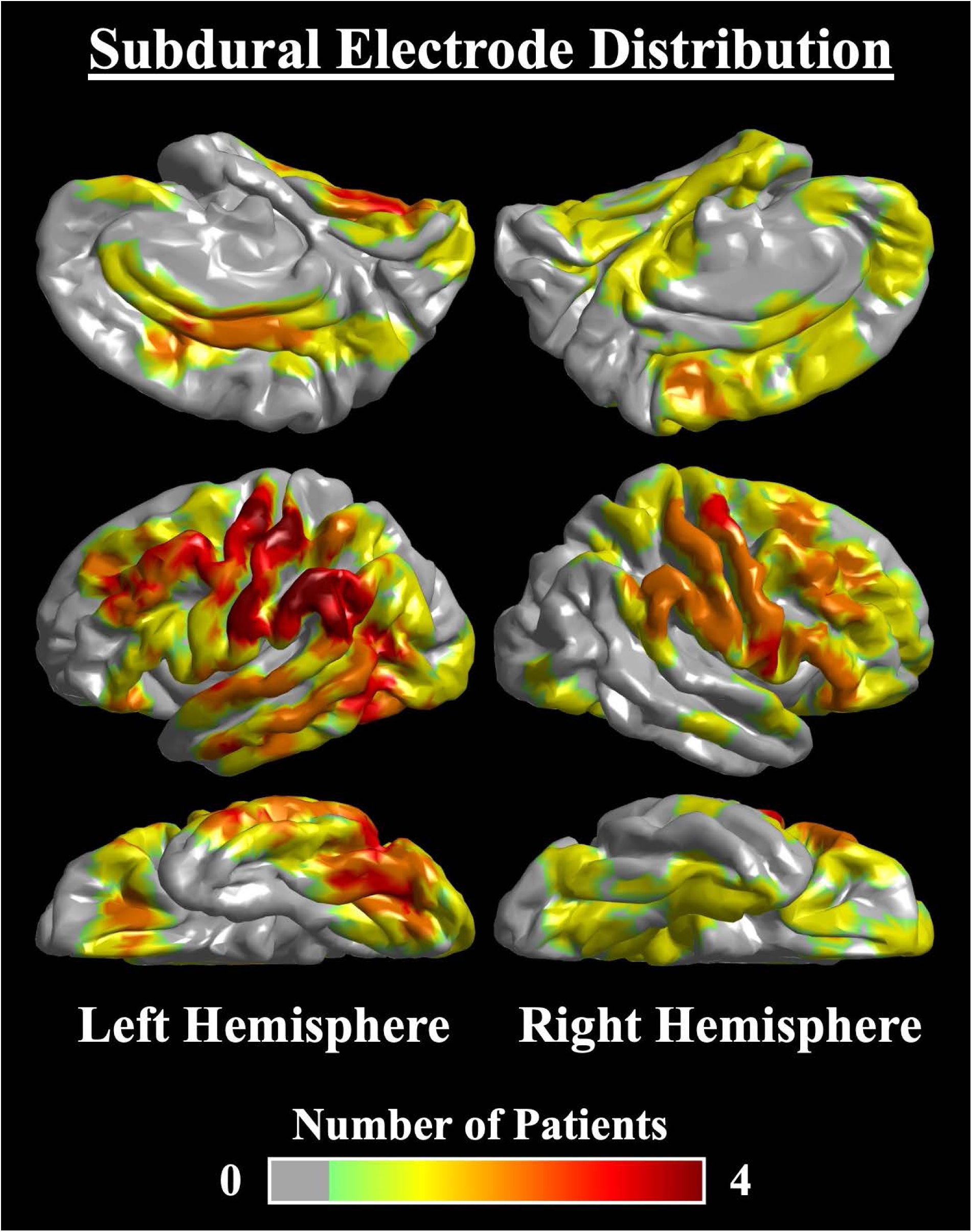
Subdural electrode distribution. Anatomical distribution of normative subdural electrodes pooled from the eight two-stage patients that underwent intraoperative electrocorticography recording at all sevoflurane stages, as well as extraoperative intracranial electroencephalography recording during slow-wave sleep (*n = 431* normative electrodes). The heatmap represents the number of patients implanted, for a given brain region.

### Dynamic tractography

Dynamic tractography was computed, as done previously, to better understand the cortico-cortical white matter pathways facilitating propagation of sevoflurane-activated spike-and-waves, approximated by delta-HFO PAC, as well as HFO.^66,69,84–87^ For each anesthetic stage, DSI Studio software (https://dsi-studio.labsolver.org/) visualized DWI white matter tracts directly connecting cortical sites showing significant co-augmentation (i.e., at least three standard deviations over the slow wave sleep mean) of the ECoG/iEEG biomarkers. Electrode positions were converted to regions on the Lausanne Brain Atlas^88^ and overlayed on the Human Connectome Project (HCP) whole-brain, white matter template.^89^ The template was made using whole-brain seeding with the following parameters: tracking threshold of 0.7, angular threshold of 70 degrees, and step-size of 0.3 mm. The open-source HCP dataset includes averaged diffusion MRI scans from approximately 1,000 participants. Diffusion MRI data was acquired using a Siemens 3T Skyra scanner with a 2D spin-echo single-shot multiband EPI sequence, a multiband factor of 3, and a monopolar gradient pulse. The spatial resolution was 1.25 mm isotropic.^89^

### Statistical analysis

#### Relationship between anesthetic stage and ECoG biomarkers

A linear mixed model determined the normative biomarker response to increasing sevoflurane concentration. Electrodes from all individual patients were pooled for analysis. The dependent variable was biomarker value. Fixed effect predictors included: [1] age, [2] biological sex, [3] sampled hemisphere, [4] presence of MRI lesion, [5] presence of daily seizures, [6] number of oral anti-seizure medications taken immediately prior to the ECoG recording, [7] one-versus two-stage surgery, and [8] anesthetic stage. Random effect predictors included patient and intercept. A fixed effect predictor greater than zero suggests that ECoG biomarkers increased as a function of sevoflurane concentration, and vice versa. This analysis was repeated for three conditions (isoflurane versus slow-wave sleep, isoflurane versus sevoflurane, and slow wave sleep versus sevoflurane), and two electrophysiology biomarkers (delta-HFO PAC and HFO EC). Thus, we employed a Bonferroni correction for 6 comparisons, taking a two-sided, corrected p-value of less than 0.05 as significant.

#### Fidelity of sevoflurane-activated ECoG biomarkers for intraoperative characterization of epileptogenicity

A binary logistic mixed model was used to determine if relatively more augmentation of the ECoG/iEEG biomarkers could characterize the epileptogenic status of electrode sites. Electrodes from all individual patients were pooled for analysis. The dependent variable was epileptogenic status (yes/no). Fixed effect predictors included: [1] age, [2] biological sex, [3] sampled hemisphere, [4] presence of MRI lesion, [5] presence of daily seizures, [6] number of anti-seizure medications, [7] one-versus two-stage surgery, and [8] biomarker value. Random effect predictors included patient and intercept. An odds ratio greater than one suggests that higher sevoflurane-activated ECoG/iEEG biomarkers were associated with an increased likelihood of characterizing a given site as epileptogenic. This analysis was repeated for the data z-scored via both the isoflurane and slow wave sleep controls, two electrophysiology epilepsy biomarkers (delta-HFO PAC and HFO EC), and ten anesthetic stages. Thus, we employed a Bonferroni correction for 40 comparisons, taking a two-sided corrected p-value of less than 0.05 as significant. Effect size of anesthetic influence on biomarker values between epileptogenic and non-epileptogenic sites was calculated using Cohen’s *d*, as done previously.^69^ For each patient and each one-minute segment of a given anesthetic stage, the Cohen’s *d* was calculated as the absolute value of: [(the average biomarker value in epileptogenic sites) minus (the average biomarker value in non-epileptogenic sites)] divided by (the standard deviation of all sites). Values for all patients and all one-minute segments of a given anesthetic stage were averaged to produce one effect size for each anesthetic stage. The same calculation was also performed for SOZ versus non-SOZ sites.

#### Seizure outcome classification

Supplementary analysis included both 19 patients who achieved ILAE-defined seizure freedom and four patients who did not achieve seizure control following surgery. The same binary logistic mixed model above was used to determine if the subtraction biomarker value could characterize seizure freedom. The subtraction biomarker value for each patient was defined as the average of all resected sites minus the average of all retained sites, as done previously.^62^ This effectively quantified the completeness of resecting delta-HFO PAC and HFO EC sites. Higher subtraction values reflect more complete resection.

## Results

### Normative sevoflurane distribution of delta-HFO phase-amplitude coupling

We first assessed the normative response of interictal, intraoperative delta-HFO PAC (rated by modulation index) to step-wise increasing sevoflurane. For the isoflurane control, linear mixed model analysis demonstrated that normative delta-HFO PAC values increased as a function of sevoflurane concentration (Fig. 3A; fixed effect estimate = 0.007; t-value = 18.566; uncorrected p-value = 2.24E-74; df = 4,825). The same trend was seen in the slow wave sleep control (Fig. 3B; fixed effect estimate = 0.006; t-value = 16.407; uncorrected p-value = 8.64E-59; df = 4,478). Linear mixed model fitting performance was assessed via Q-Q plots of residuals for both controls (Supplementary Fig. 1). Similar calculation did not show a difference between normative isoflurane (1 vol%) and slow wave sleep (fixed effect estimate = −0.002; t-value = −0.670; uncorrected p-value = 0.503; df = 1,240). Figure 3C visualizes the cortical surface distribution of delta-HFO PAC at each anesthetic stage, for eight patients that received extraoperative iEEG recording plus every intraoperative stage (*n = 431* normative sites). Prominent posterior delta-HFO PAC enhancement during the slow wave sleep condition shifted to sensorimotor prominence by sevoflurane (4 vol%). Dynamic tractography suggested that the significant sevoflurane-induced delta-HFO PAC co-augmentation between Rolandic areas was supported by inter-hemispheric callosal fibers (Fig. 3D).

**Figure 3.**
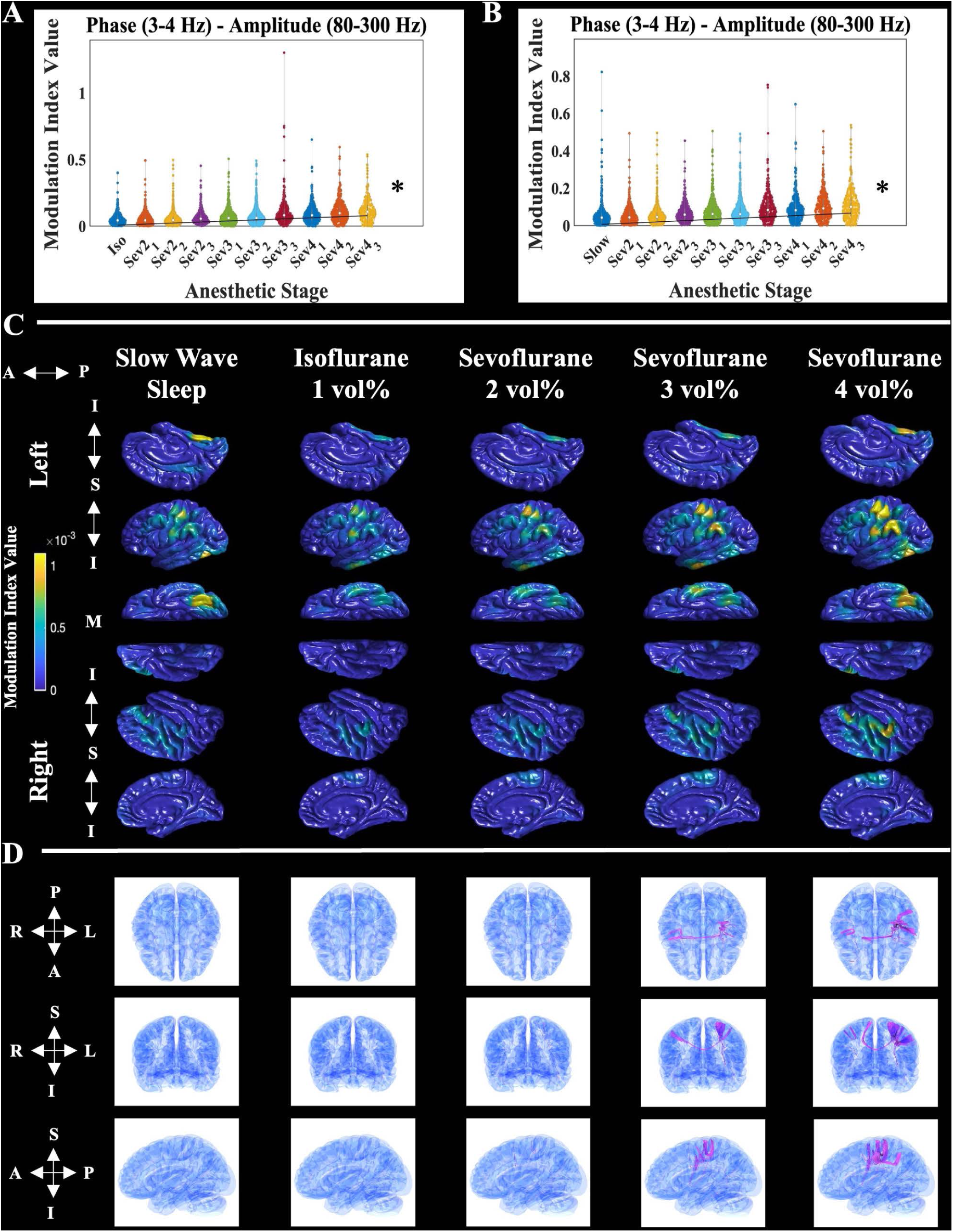
Normative sevoflurane distribution of delta and high-frequency oscillation phase-amplitude coupling. Mathematical distribution of ECoG/iEEG-defined delta (3-4 Hz) and HFO (80-300 Hz) phase-amplitude coupling (delta-HFO PAC) – rated by modulation index -from all pooled normative electrode sites, at each anesthetic stage. The left panel **(A)** shows delta-HFO PAC data from one-stage and two-stage patients (iso; *n = 963* normative electrode sites) and a linear mixed model that uses the isoflurane period as the reference. The right panel **(B)** shows delta-HFO PAC data from two-stage patients (slow; *n = 816* normative electrode sites) and a linear mixed model that uses the slow wave sleep period as the reference. In addition, ‘Sev2_1’ denotes the first minute of sevoflurane at a concentration of two-volume-percent, and so on. The black trend line anchored at the origin depicts the relationship between delta-HFO PAC and anesthetic stage: determined by linear mixed model analysis. The asterisks represent a significant effect (Bonferroni corrected p < 0.05) of anesthetic stage on biomarker value, via linear mixed model analysis. (**C)** Anatomical distribution of normative delta-HFO PAC interpolated onto the FreeSurfer average cortical surface. The data represents normative electrode sites pooled from the eight patients who underwent both extraoperative iEEG recording during slow wave sleep, as well as intraoperative ECoG at every anesthetic stage (*n = 431* normative electrode sites). Each column represents a different anesthetic stage. The left and right hemispheres are grouped in the first and last three rows, respectively. Within each hemisphere group, single rows represent different cortical surfaces. For the left hemisphere: top row-sagittal-medial, middle row-sagittal-lateral, and bottom row-axial-inferior. The orientation row order is opposite for the right hemisphere. The anterior – posterior (‘A ←→ P’) orientation is listed for all views. In addition, the inferior and superior orientations for the sagittal images are depicted (‘I ←→S ←→ I’), as well as the medial (‘M’) marker for the inferior-axial view. Hotter colors represent sites with relatively higher ECoG biomarker values and vice versa. The spatial coverage of intracranial electrodes is presented in Figure 2. In areas without intracranial electrode coverage, the absence of biomarker values should be interpreted as a lack of intracranial EEG signal sampling. (**D)** Dynamic tractography depicts diffusion weighted-imaging (DWI) white matter streamlines connecting cortical sites with significantly elevated delta-HFO PAC (i.e., greater than three standard deviations above the slow wave sleep mean). Each column represents a different anesthetic stage, and each row, from top to bottom, shows a different brain view: axial, coronal, and sagittal, respectively. The orientation for each view is included to the left of the corresponding row. A = anterior; extraoperative iEEG = extraoperative intracranial electroencephalography; I = inferior; intraoperative ECoG = intraoperative electrocorticography; HFO = high-frequency oscillation; L = left; M = medial; P = posterior; PAC = phase-amplitude coupling; R = right; S = superior.

### Normative sevoflurane distribution of HFO effective connectivity

We next determined the normative response of interictal HFO EC (rated by transfer entropy) to step-wise increasing sevoflurane. Linear mixed model analysis suggested that HFO EC increased as a function of sevoflurane concentration, for both controls (Fig. 4A; isoflurane control: fixed effect estimate = 6.076E-5; t-value = 8.248; uncorrected p-value = 2.15E-16; df = 4,008 | Fig. 4B; slow wave sleep control: fixed effect estimate = 4.107E-5; t-value = 6.806; uncorrected p-value = 1.14E-11; df = 4,419). Linear mixed model fitting performance was assessed via Q-Q plots of residuals for both controls (Supplementary Fig. 2). Separating afferent and efferent transfer entropy suggested they behaved the same as combined (Supplementary Table 2). There was not a significant difference between normative isoflurane (1 vol%) and slow wave sleep (fixed effect estimate = 8.207E-5; t-value = 1.199; uncorrected p-value = 0.231; df = 1,239). Interpolation of normative sites (*n = 431* electrodes) onto an average cortical surface qualitatively showed sevoflurane-activated HFO EC in frontal and parietal areas (Fig. 4C). Dynamic tractography traced prominent intra-hemispheric connections between significant sevoflurane-activated cortical HFO EC foci in frontal and parietal association regions, via the superior longitudinal fasciculus (Fig. 4D).

**Figure 4.**
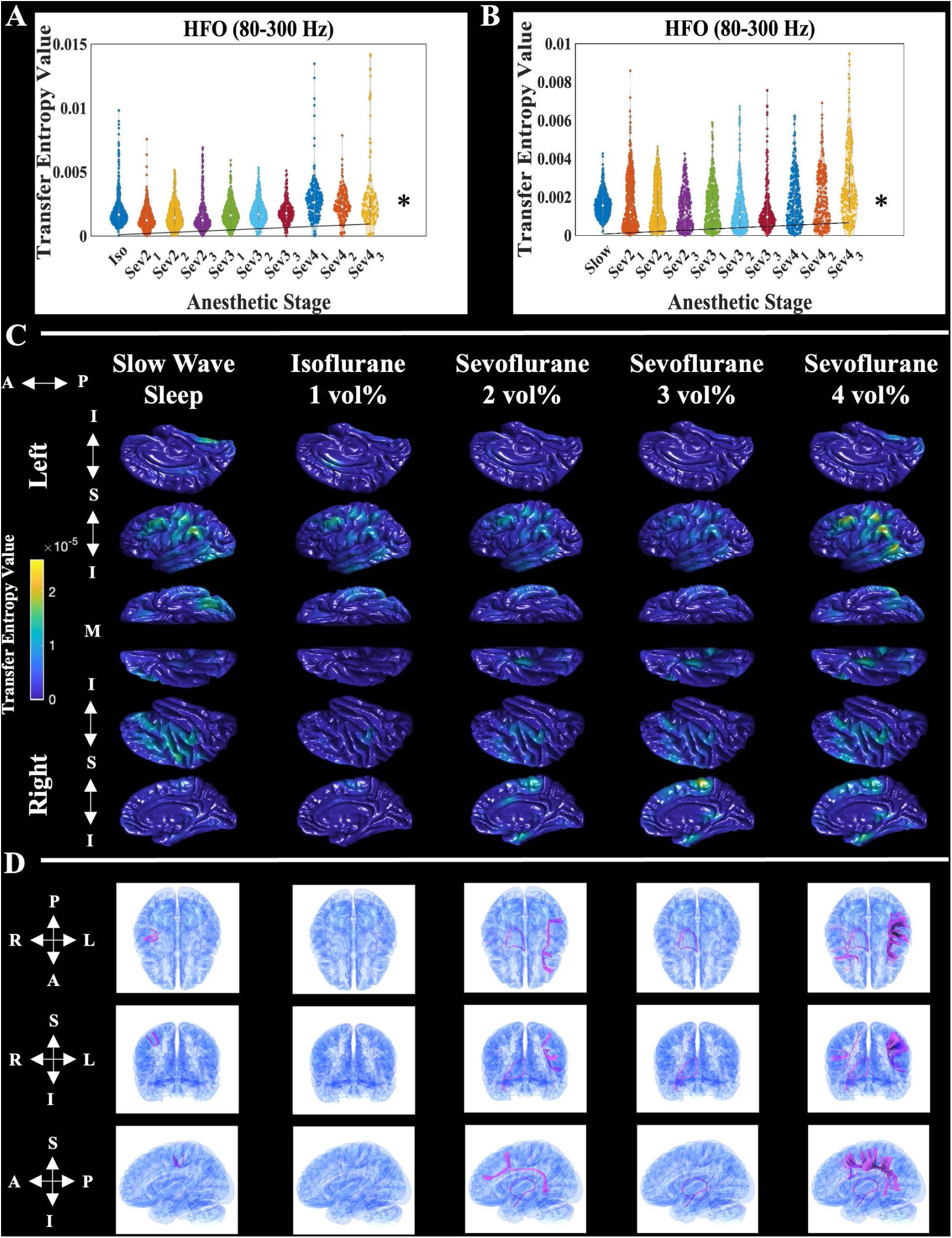
Normative sevoflurane distribution of high-frequency oscillation effective connectivity. Mathematical distribution of high-frequency oscillation (80-300 Hz) effective connectivity – rated by transfer entropy -from all pooled normative electrode sites, at each anesthetic stage. The left panel **(A)** shows HFO EC data from one-stage and two-stage patients derived from a spectral amplitude time series z-scored in relation to the mean and standard deviation of the the isoflurane control (iso; *n = 963* normative electrode sites) and a linear mixed model that uses the isoflurane period as the reference. The right panel **(B)** shows HFO EC data from two-stage patients derived from a spectral amplitude time series z-scored in relation to the mean and standard deviation of the the slow wave sleep control (slow; *n = 816* normative electrode sites) and a linear mixed model that uses the slow wave sleep period as the reference. In addition, ‘Sev2_1’ denotes the first minute of sevoflurane at a concentration of two-volume-percent, and so on. The black trend line anchored at the origin depicts the relationship between HFO effective connectivity and anesthetic stage: determined by linear mixed model analysis. The asterisks represent a significant effect (Bonferroni corrected p-values < 0.05) of anesthetic stage on biomarker value, via linear mixed model analysis. (**C)** Anatomical distribution of normative HFO EC interpolated onto the FreeSurfer average cortical surface. The data represents normative electrode sites pooled from the eight patients who underwent both extraoperative iEEG recording during slow wave sleep, as well as intraoperative ECoG at every anesthetic stage (*n = 431* normative electrode sites). Each column represents a different anesthetic stage. The left and right hemispheres are grouped in the first and last three rows, respectively. Within each hemisphere group, single rows represent different cortical surfaces. For the left hemisphere: top row-sagittal-medial, middle row-sagittal-lateral, and bottom row-axial-inferior. The orientation row order is opposite for the right hemisphere. The anterior – posterior (‘A ←→ P’) orientation is listed for all views. In addition, the inferior and superior orientations for the sagittal images are depicted (‘I ←→S ←→ I’), as well as the medial (‘M’) marker for the inferior-axial view. Hotter colors represent sites with relatively higher biomarker values and vice versa. The spatial coverage of intracranial electrodes is presented in Figure 2. In areas without intracranial electrode coverage, the absence of biomarker values should be interpreted as a lack of intracranial EEG signal sampling. (**D)** Dynamic tractography depicts diffusion weighted-imaging (DWI) white matter streamlines connecting cortical sites with significantly elevated HFO EC (i.e., greater than three standard deviations above the slow wave sleep mean). Each column represents a different anesthetic stage, and each row, from top to bottom, shows a different brain view: axial, coronal, and sagittal, respectively. The orientation for each view is included to the left of the corresponding row. A = anterior; DWI = diffusion weighted imaging; EC = effective connectivity; extraoperative iEEG = extraoperative intracranial electroencephalography; I = inferior; intraoperative ECoG = intraoperative electrocorticography; HFO = high-frequency oscillation; L = left; M = medial; P = posterior; R = right; S = superior.

### Normative sevoflurane distribution of delta effective connectivity

We created a supplementary normative map of interictal, sevoflurane-activated delta EC (rated by transfer entropy). Linear mixed model analysis showed that sevoflurane increased delta EC, regardless of control (Supplementary Fig. 3A-B and Supplementary Table 3; significance with Bonferroni corrected p-values < 0.05). Linear mixed model fitting performance was assessed via Q-Q plots of residuals for both controls (Supplementary Fig. 4). Combined, afferent, and efferent delta transfer entropy all responded to sevoflurane in the same manner (Supplementary Table 3; significance with Bonferroni corrected p-values < 0.05). Compared to slow wave sleep, isoflurane increased delta EC (Supplementary Table 3; significance with Bonferroni corrected p-values < 0.05). Normative sites (*n = 431* electrodes) pooled on the FreeSurfer average cortical surface qualitatively showed prominent delta EC activations in frontal and parietal association areas, along with smaller foci in the sensorimotor strips (Supplementary Fig. 3C). Dynamic tractography suggested that the significant normative delta EC hotspots were connected via both inter-hemispheric callosal fibers and intra-hemispheric superior longitudinal fasciculi (Supplementary Fig. 3D).

### Characterizing epileptogenicity via intraoperative ECoG biomarkers

In the next experiment, we pooled all electrode sites, regardless of normative status. Binary logistic mixed model analysis suggested that relatively higher intraoperative delta-HFO PAC values were associated with an increased likelihood of characterizing a given site as epileptogenic, at every anesthetic stage (Fig. 5A-B and Table 2). A double dissociation was found between HFO and delta EC values. Compared to non-epileptogenic sites, greater HFO EC (Fig. 5C-D and Table 3) and lower delta EC (Supplementary Fig. 5 and Supplementary Table 4) were associated with epileptogenic electrode sites at sevoflurane (3-4 vol%) and sevoflurane (2-4 vol%), respectively. Greater delta-HFO PAC and HFO EC in epileptogenic sites was also noted during isoflurane (1 vol%) and slow wave sleep stages. However, Cohen’s *d* calculation (Supplementary Table 5) showed that this effect was higher during sevoflurane (3-4 vol%). Since epileptogenicity can’t be determined until the seizure outcome is known, we also conducted a parallel supplementary analysis characterizing the SOZ, which is estimated prior to resection surgery. The binary logistic mixed model and Cohen’s *d* analysis showed the same relationship, especially at sevoflurane (3-4 vol%): higher delta-HFO PAC and HFO EC but lower delta EC augmentation characterized SOZ sites (Supplementary Fig. 6 and Supplementary Tables 6-9).

**Figure 5.**
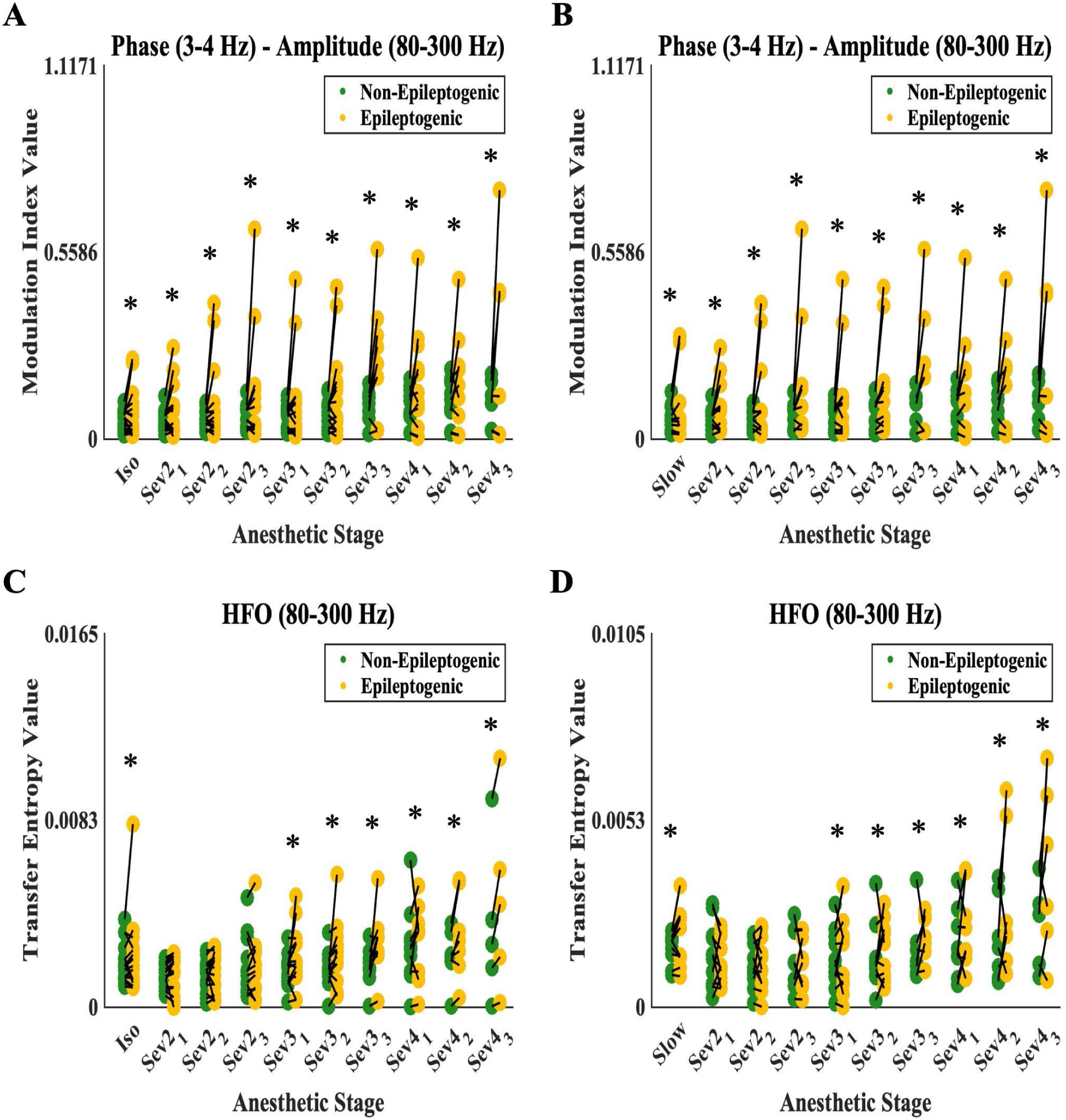
Sevoflurane-activated ECoG biomarkers characterize epileptogenicity. Delta (3-4 Hz) and high-frequency oscillation (80-300 Hz) phase-amplitude coupling (delta-HFO PAC rated by modulation index), values from one-stage and two-stage patients in the **(A)** isoflurane control distribution (*n = 1,608* total electrode sites) and from two-stage patients in the **(B)** slow wave sleep control distribution (*n = 1,344* total electrode sites), at each anesthetic stage. For a given patient, the green and yellow dots represent the average value of all non-epileptogenic and epileptogenic sites, respectively. In each anesthetic stage, the black lines connect a pair of green-yellow dots from the same patient. The asterisks denote binary logistic mixed model significance (Bonferroni corrected *p < 0.05*) for characterizing epileptogenicity via iEEG/ECoG biomarker levels. On the x-axis, ‘Sev2_1’ denotes the first minute of sevoflurane at a concentration of two-volume-percent, and so on. The same analysis is repeated for HFO effective connectivity (rated by transfer entropy). Panel **(C)** shows HFO EC values from one-stage and two-stage patients derived from a spectral amplitude time series z-scored against the mean and standard deviation of the isoflurane control (*n = 1,608* total electrode sites). Panel **(D)** shows HFO EC values from two-stage patients derived from a spectral amplitude time series z-scored against the mean and standard deviation of the slow wave sleep control (*n = 1,344* total electrode sites). ‘Iso’ refers to the isoflurane period, while ‘slow’ refers to the slow wave sleep period. EC = effective connectivity; Extraoperative iEEG = extraoperative intracranial electroencephalography; HFO = high-frequency oscillation; intraoperative ECoG = intraoperative electrocorticography; PAC = phase-amplitude coupling.

### Intraoperative prediction of seizure freedom

A final supplementary analysis sought to determine if the completeness of resecting regions displaying high values of sevoflurane-activated delta-HFO PAC and/or EC could predict post-operative ILAE class-I seizure freedom. For each patient, subtracting the average ECoG/iEEG biomarker value of all retained sites from the average of all resected sites served as a proxy for resection completeness. However, the subtraction values did not significantly predict seizure freedom (Supplementary Tables 10-12).

## Discussion

### Significance and innovation

To our knowledge, this is the first study to depict the normative distribution of ECoG/iEEG-based intraoperative delta-HFO PAC and HFO EC, at step-wise increasing concentrations of sevoflurane anesthesia. The novelty of this work includes: [1] beginning to illuminate the normative cortical distribution of ECoG/iEEG-based delta-HFO PAC, HFO EC, and delta EC, at various sevoflurane, isoflurane, and slow wave sleep stages; [2] using dynamic tractography to delineate cortico-cortical white matter pathways potentially facilitating the propagation of sevoflurane-induced delta-HFO PAC and HFO EC across normative areas; and [3] showing the fidelity of sevoflurane-activated delta-HFO PAC, HFO EC, and delta EC for intraoperative characterization of epileptogenic and SOZ sites.

Relatively few studies have investigated the effects of sevoflurane on HFO signaling in patients with focal epilepsy. Previous studies reported that the occurrence rates of spikes and HFO increased most prominently in the SOZ under sevoflurane.^50,51^ The present findings also align with our preliminary investigation, which analyzed intraoperative ECoG data from a separate cohort of eight patients with drug-resistant focal epilepsy. Data from both cohorts independently suggests that sevoflurane generally elevates delta-HFO phase-amplitude coupling and HFO effective connectivity, with the greatest increases observed in epileptogenic sites.^68,69^ Moreover, these studies demonstrated a double dissociation, wherein epileptogenic regions exhibit relatively higher delta-HFO PAC and HFO effective connectivity but lower delta effective connectivity augmentation under sevoflurane. Slow waves are thought to reflect neural inhibition;^90–92^ whereas, HFO coupled with spike-and-wave discharges are attributed to neural excitation.^93,94^ This supports the notion that sites experiencing relatively less inhibitory delta influence concurrently display greater levels of HFO. Another recent study by Uda et al. (2024) showed that the highest degree of sevoflurane-enhanced HFO and spike rates were in SOZ sites.^72^ While Uda et al. (2024) examined the occurrence rates of HFO events in single channels, the present study analyzed HFO coupling with slow waves and network properties, specifically delta-HFO phase-amplitude coupling and effective connectivity. In addition, the current investigation included dynamic tractography to delineate possible white matter pathways facilitating HFO propagation under sevoflurane. The results of these analyses infer that the ability to characterize epileptogenicity was optimized between sevoflurane 3-4 vol%.^68,69,72,95^

Numerous studies suggest that HFO serve as a compelling biomarker of epilepsy, with their removal reportedly associated with postoperative seizure freedom.^40–69,96–98^ Clinical investigations have also implicated elevated delta-HFO phase-amplitude coupling in identifying seizure foci.^57–59,62–66,69,99–101^ However, it is established that physiologic HFO exist in the brain and are thought to play a role in memory consolidation.^73,102–104^ Similarly, physiologic phase-amplitude coupling is suggested to integrate distant spatial signals into singular cognitive events.^67,105–106^ When considering these ECoG features as epilepsy biomarkers, distinguishing physiologic activity from genuinely pathological signals is crucial. This distinction is particularly relevant given that sevoflurane is known to induce epileptiform EEG signals and seizure-like movements even in non-epileptic patients.^29–30^ Focusing on HFO network dynamics may help address this challenge, as epilepsy is associated with abnormal network connectivity.^69,107–112^ Our results suggest that while sevoflurane enhances HFO effective connectivity even in normative sites, the most pronounced augmentation occurs in the epileptogenic and seizure onset zones. This aligns with studies demonstrating that distinct spike and HFO propagation patterns characterize epileptogenic brain networks.^69,113–115^ Specifically, Tamilia et al. (2021) and others have suggested that within pathological brain networks, the onset location of HFO and spike propagation trains is most indicative of the epileptogenic zone.^116–119^ Our dynamic tractography results attempt to provide an anatomical framework for these neural propagations, albeit within normative sites under sevoflurane. Previous clinical iEEG studies have implicated white matter as a conduit for epileptiform signal propagation.^69,85,120–123^ Consequently, these fibers may serve as critical targets for surgical disconnection to prevent pathological discharges from reaching regions responsible for clinical manifestations.

Our novel normative sevoflurane brain maps provide a preliminary reference to improve distinguishing off-target anesthetic-related effects from genuine epileptogenic augmentation. While more investigation is needed, delta-HFO phase-amplitude coupling (rated by modulation index) and HFO effective connectivity (rated by transfer entropy), represent exciting objective intraoperative ECoG epilepsy biomarkers. Using sevoflurane to rapidly and reversibly augment these electrophysiological metrics during surgery could help reduce a severe diagnostic burden by mitigating the need for intracranial electrode implants and extraoperative iEEG recording. This would optimize treatment cost effectiveness, while maintaining post-operative seizure outcomes comparable with conventional methods. Even in cases when two-stage surgery is still necessary, sevoflurane-based guidance could improve intracranial electrode placement, although future studies are needed to confirm this point.

### Possible mechanisms

While the exact mechanism of how sevoflurane enhances epileptiform HFO signal generation remains unclear, we hypothesize the following working model. It is accepted that sevoflurane potentiates inhibitory GABAergic neurotransmission.^124–127^ This hyperpolarizing drive disinhibits T-type calcium channels in thalamocortical relay neurons, which enter burst firing mode. Bursting in the thalamus entrains cortical synchronization and, via corticothalamic projections, sets up a feedback resonance with the thalamus.^128–133^ Cortical interneurons organize the thalamocortical bursts into delta waves.^128–133^ However, extensive sevoflurane exposure is known to overwhelm the cellular GABAergic machinery responsible for coordinating cortical activity, and this impaired inhibition leads to asynchronous neuronal firing.^127,134–136^ HFO might then emerge from the combination of malfunctioning perisomatic inhibition plus large waves of sevoflurane-driven bursting activity, leading to irregular summation of asynchronous cortical firing.^91^ Since epileptic tissue is thought to have disrupted inhibition,^137–138^ it would inherently be more sensitive to such effects. This phenomenon might also help explain the double dissociation between higher delta-HFO PAC and HFO EC but lower delta EC, in epileptogenic and SOZ sites. Sevoflurane (3-4 vol%) may have paradoxically blunted delta activity, disinhibiting HFO and spike-and-wave propagation, even in normative brain regions. This model is further supported by the findings that [1] the GABA_A_ antagonist bicuculline can cause epileptiform discharges and [2] penicillin is believed to operate via a similar mechanism to spur EEG spikes.^134,139^ We believe the present study begs basic science questions on how sevoflurane alters neural signaling. Better understanding of the cellular and circuit mechanisms generating HFO and delta waves could provide avenues to improve the system for optimal clinical use.

### Limitations and future directions

There were several limitations to the present work that should be addressed. One criticism is the small number of patients used in this study. Thus, it is possible that the qualitative signal distribution displayed on the brain maps may be partially skewed toward the anatomical distribution of implanted electrodes, in these few patients. Ethical concerns constrain invasive sampling only to implicated brain regions. Increasing the number of subjects would certainly improve resolution. However, we were limited to the number of patients that fit inclusion criteria. The mixed model approach helps overcome this issue because: [1] it allows each individual electrode site to count toward the *n*-number, and [2] it allows pooling of electrodes across all patients. Since ‘patient’ was included as a random-effects variable in the mixed models, baseline differences across subjects were inherently accounted for. That allowed us to pool all electrode sites together for the analysis. For example, the isoflurane control period included *n = 1,608* electrode sites; meaning that with a power of 0.8 and an alpha = 0.05 we were able to detect a small effect size of around 0.01. Thus, our models have adequate statistical power to assess the impact of sevoflurane on the ECoG/iEEG biomarkers.

However, our subtraction models applied to 23 patients did not demonstrate that ECoG/iEEG biomarkers could distinguish between patients who achieved post-operative seizure freedom and those who did not. We computed subtraction as the difference between the average of all retained and resected sites, treating this measure as an estimate of the completeness of biomarker site resection. A previous study from our group involving over 100 patients showed that subtraction biomarker values during slow-wave sleep classified patients with ILAE class I seizure outcomes.^62^ An alternative analytic approach may better quantify the completeness of resecting high-value biomarker sites. Future studies may benefit from normalizing subtraction values, as performed in a study of 109 patients,^55^ to mitigate baseline differences across patients before statistical analysis. The failure to demonstrate the predictive utility of ECoG/iEEG biomarkers may be attributed to the small sample size. The present analysis included only 4 poor-outcome patients and 19 ILAE class I seizure-free patients. With a sample size of 23 and α = 0.05, statistical power was only 0.32 to detect a large effect size (0.8). Increasing the sample size to 80 would be necessary to achieve sufficient power to detect this effect. Another criticism involves pooling one- and two-stage surgery patients for analysis. We accounted for this potential confounder in our statistical models, which failed to find any significant effect of surgical staging on the outcomes. Also, both groups received identical intraoperative ECoG recording paradigms in the presence of step-wise increasing sevoflurane. The same may be said for pooling patients aged 4-18 years because the brain undergoes dramatic development during that time. The anatomical distribution of physiologic delta-HFO PAC shifts as children grow.^66^ The best way to account for these dynamic cerebral changes would be to stratify our analysis based on patient age. However, we did not have the appropriate sample size to conduct such analysis. We, instead, incorporated age as a fixed effect factor in our mixed model statistics, as done previously^62,68–69^. Our analysis in this study likewise failed to find a significant effect of age on outcomes. At least within the bounds of the studies reported here, treatment plan and age did not affect outcome.

Volume conduction is another potential issue impacting interpretation of delta-HFO PAC and EC. Mercier et al. (2022) published guidelines and good practices for analysis of human iEEG.^140^ They suggest that local bipolar referencing is the optimal referencing scheme for ECoG/iEEG-derived HFO. This technique mitigates undesired effects such as volume conduction and other artifacts. We employed bipolar referencing in the current study to account for such effects, but it must be considered that this technique does not fully eliminate these factors. Finally, not all electrodes were uniformly included for each anesthetic stage due to either electrographic artifacts and/or, in limited cases, physiologic factors prompting the anesthesiologist to cease increasing sevoflurane. The mixed model approach sufficiently handled this missing data.

In the future, we want to consider prospective studies to: [1] definitively assess whether adding sevoflurane-guidance to intracranial electrode implantation leads to better post-operative seizure control, and [2] to determine if bolstering standard spike-based visualization with delta-HFO PAC and HFO EC can improve the chances of post-operative seizure freedom.

## Data availability

All data used in this study is available upon reasonable request to the corresponding author. Transfer entropy code is available at GitHub: https://github.com/efire11/Transfer-Entropy.git

## Supplementary material

Supplementary material is available at *Brain Communications* online.

## References

1. Laguitton V, Desnous B, Lépine A, et al. Intellectual outcome from 1 to 5 years after epilepsy surgery in 81 children and adolescents: a longitudinal study. Seizure. 2021;91:384–392.

2. Morningstar M, French RC, Mattson WI, Englot DJ, Nelson EE. Social brain networks: resting-state and task-based connectivity in youth with and without epilepsy. Neuropsychologia. 2021;157:107882.

3. Sultana B, Panzini MA, Veilleux Carpentier A, et al. Incidence and prevalence of drug-resistant epilepsy: a systematic review and meta-analysis. Neurology. 2021;96:805–817.

4. Wiebe S, Blume WT, Girvin JP, Eliasziw M. Effectiveness and Efficacy of Surgery for Temporal Lobe Epilepsy Study Group. A randomized, controlled trial of surgery for temporal-lobe epilepsy. N Engl J Med. 2001;345:311–318.

5. Dwivedi R, Ramanujam B, Chandra PS, et al. Surgery for drug-resistant epilepsy in children. N Engl J Med. 2017;377:1639–1647.

6. Widjaja E, Jain P, Demoe L, Guttmann A, Tomlinson G, Sander B. Seizure outcome of pediatric epilepsy surgery: systematic review and meta-analyses. Neurology. 2020;94:311–321.

7. Hader WJ, Mackay M, Otsubo H, et al. Cortical dysplastic lesions in children with intractable epilepsy: role of complete resection. J Neurosurg. 2004;100:110–117.

8. Lüders HO, Najm I, Nair D, Widdess-Walsh P, Bingman W. The epileptogenic zone: general principles. Epileptic Disord. 2006;8:S1–S9.

9. Asano E, Juhász C, Shah A, Sood S, Chugani HT. Role of subdural electrocorticography in prediction of long-term seizure outcome in epilepsy surgery. Brain. 2009;132:1038–1047.

10. Jayakar P, Gotman J, Harvey AS, et al. Diagnostic utility of invasive EEG for epilepsy surgery: indications, modalities, and techniques. Epilepsia. 2016;57:1735–1747.

11. Mullin JP, Shriver M, Alomar S, et al. Is SEEG safe? A systematic review and meta-analysis of stereo-electroencephalography-related complications. Epilepsia. 2016;57:386–401.

12. Uribe-Cardenas R, Boyke AE, Schwarz JT, et al. Utility of invasive electroencephalography in children 3 years old and younger with refractory epilepsy. J Neurosurg Pediatr. 2020;26:1–6.

13. Roth J, Constantini S, Ekstein M, et al. Epilepsy surgery in infants up to 3 months of age: safety, feasibility, and outcomes: a multicenter, multinational study. Epilepsia. 2021;62:1897–1906.

14. Gavvala J, Zafar M, Sinha SR, Kalamangalam G, Schuele S, American SEEG Consortium, supported by The American Clinical Neurophysiology Society. Stereotactic EEG practices: a survey of United States tertiary referral epilepsy centers. J Clin Neurophysiol. 2022;39:474–480.

15. Bernabei JM, Li A, Revell AY, et al. Quantitative approaches to guide epilepsy surgery from intracranial EEG. Brain. 2023;146:2248–2258.

16. Hader WJ, Tellez-Zenteno J, Metcalfe A, et al. Complications of epilepsy surgery: a systematic review of focal surgical resections and invasive EEG monitoring. Epilepsia. 2013;54:840–847.

17. Belohlavkova A, Jezdik P, Jahodova A, et al. Evolution of pediatric epilepsy surgery program over 2000-2017: improvement of care? Eur J Paediatr Neurol. 2019;23:456–465.

18. Krsek P, Jahodova A, Kyncl M, et al. Predictors of seizure-free outcome after epilepsy surgery for pediatric tuberous sclerosis complex. Epilepsia. 2013;54:1913–1921.

19. Bansal S, Kim AJ, Berg AT, et al. Seizure outcomes in children following electrocorticography-guided single-stage surgical resection. Pediatr Neurol. 2017;71:35–42.

20. Jing J, Herlopian A, Karakis I, et al. Interrater reliability of experts in identifying interictal epileptiform discharges in electroencephalograms. JAMA Neurol. 2020;77:49–57.

21. Hu DK, Rana M, Adams DJ, et al. Interrater reliability of interictal EEG waveforms in Lennox-Gastaut Syndrome. Epilepsia Open. 2024;9:176–186.

22. Gordon E, Widen L. General anesthesia with halothane for surgical interventions and electrocorticography in cases of focal epilepsy. Acta Anaesthesiol Scand. 1962;6:13–28.

23. Sato Y, Sato K, Shamoto H, Kato M, Yoshimoto T. Effect of nitrous oxide on spike activity during epilepsy surgery. Acta Neurochir (Wien). 2001;143:1213–1216.

24. Asano E, Benedek K, Shah A, et al. Is intraoperative electrocorticography reliable in children with intractable neocortical epilepsy? Epilepsia. 2004;45:1091–1099.

25. Kacar Bayram A, Yan Q, Isitan C, Rao S, Spencer DD, Alkawadri R. Effect of anesthesia on electrocorticography for localization of epileptic focus: literature review and future directions. Epilepsy Behav. 2021;118:107902.

26. Komatsu H, Taie S, Endo S, et al. Electrical seizures during sevoflurane anesthesia in two pediatric patients with epilepsy. Anesthesiology. 1994;81:1535–1537.

27. Woodforth IJ, Hicks RG, Crawford MR, Stephen JPH, Burke DJ. Electroencephalographic evidence of seizure activity under deep sevoflurane anesthesia in a nonepileptic patient. Anesthesiology. 1997;87:1579–1582.

28. Iijima T, Nakamura Z, Iwao Y, Sankawa H. The epileptogenic properties of the volatile anesthetics sevoflurane and isoflurane in patients with epilepsy. Anesth Analg. 2000;91:989–995.

29. Schultz A, Schultz B, Grouven U, Korsch G. Epileptiform activity in the EEGs of two nonepileptic children under sevoflurane anaesthesia. Anaesth Intensive Care. 2000;28:205–207.

30. Jääskeläinen SK, Kaisti K, Suni L, Hinkka S, Scheinin H. Sevoflurane is epileptogenic in healthy subjects at surgical levels of anesthesia. Neurology. 2003;61:1073–1078.

31. Akeson J, Didriksson I. Convulsions on anaesthetic induction with sevoflurane in young children. Acta Anaesthesiol Scand. 2004;48:405–407.

32. Kurita N, Kawaguchi M, Hoshida T, Nakase H, Sakaki T, Furuya H. The effects of sevoflurane and hyperventilation on electrocorticogram spike activity in patients with refractory epilepsy. Anesth Analg. 2005;101:517–523.

33. Särkelä MO, Ermes MJ, van Gils MJ, Yli-Hankala AM, Jäntti VH, Vakkuri AP. Quantification of epileptiform electroencephalographic activity during sevoflurane mask induction. Anesthesiology. 2007;107:928–938.

34. Gibert S, Sabourdin N, Louvet N, et al. Epileptogenic effect of sevoflurane: determination of the minimal alveolar concentration of sevoflurane associated with major epileptoid signs in children. Anesthesiology. 2012;117:1253–1261.

35. Cornelissen L, Kim SE, Purdon PL, Brown EM, Berde CB. Age-dependent electroencephalogram (EEG) patterns during sevoflurane general anesthesia in infants. Elife. 2015;4:e06513.

36. Tanaka S, Oda Y, Ryokai M, et al. The effect of sevoflurane on electrocorticographic spike activity in pediatric patients with epilepsy. Paediatr Anaesth. 2017;27:409–416.

37. Cornelissen L, Kim SE, Lee JM, Brown EN, Purdon PL, Berde CB. Electroencephalographic markers of brain development during sevoflurane anaesthesia in children up to 3 years old. Br J Anaesth. 2018;120:1274–1286.

38. Stasiowski MJ, Marciniak R, Dulawa A, Krawczyk L, Jalowiecki P. Epileptiform EEG patterns during different techniques of induction of anaesthesia with sevoflurane and propofol: a randomised trial. Anaesthesiol Intensive Ther. 2019;51:21–34.

39. Edgington TL, Muco E, Maani CV, eds. Sevoflurane. StatPearls Publishing; 2023.

40. Jacobs J, LeVan P, Chander R, Hall J, Dubeau F, Gotman J. Interictal high-frequency oscillations (80-500 Hz) are an indicator of seizure onset areas independent of spikes in the human epileptic brain. Epilepsia. 2008;49:1893–1907.

41. Blanco JA, Stead M, Krieger A, et al. Data mining neocortical high-frequency oscillations in epilepsy and controls. Brain. 2011;134:2948–2959.

42. Frauscher B, Bartolomei F, Kobayashi K, et al. High-frequency oscillations: the state of clinical research. Epilepsia. 2017;58:1316–1329.

43. Van’t Klooster MA, van Klink NEC, Zweiphenning WJEM, et al. Tailoring epilepsy surgery with fast ripples in the intraoperative electrocorticogram. Ann Neurol. 2017;81:664–676.

44. Bernardo D, Nariai H, Hussain SA, et al. Visual and semi-automatic non-invasive detection of interictal fast ripples: a potential biomarker of epilepsy in children with tuberous sclerosis complex. Clin Neurophysiol. 2018;129:1458–1466.

45. Liu S, Gurses C, Sha Z, et al. Stereotyped high-frequency oscillations discriminate seizure onset zones and critical functional cortex in focal epilepsy. Brain. 2018;141:713–730.

46. Weiss SA, Berry B, Chervoneva I, et al. Visually validated semi-automatic high-frequency oscillation detection aides the delineation of epileptogenic regions during intra-operarative electrocorticography. Clin Neurophysiol. 2018;129:2089–2098.

47. Schönberger J, Huber C, Lachner-Piza D, et al. Interictal fast ripples are associated with the seizure-generating lesion in patients with dual pathology. Front Neurol. 2020;11:573975.

48. Guth TA, Kunz L, Brandt A, et al. Interictal spikes with and without high-frequency oscillations have different single-neuron correlates. Brain. 2021;144:3078–3088.

49. Lin J, Smith GC, Gliske SV, Zochowski M, Shedden K, Stacey WC. High frequency oscillation network dynamics predict outcome in non-palliative epilepsy surgery. Brain Commun. 2024;6:fcae032.

50. Orihara A, Hara K, Hara S, et al. Effects of sevoflurane anesthesia on intraoperative high-frequency oscillations in patients with temporal lobe epilepsy. Seizure. 2020;82:44–49.

51. Orihara A, Inaji M, Fujii S, Fujimoto SH, Hara K, Maehara T. Validity of intraoperative ECoG in the parahippocampal gyrus as an indicator of hippocampal epileptogenicity. Epilepsy Res. 2022;184:106950.

52. Zweiphenning W, van’t Klooster MA, van Klink NEC, et al. Intraoperative electrocorticography using high-frequency oscillations or spikes to tailor epilepsy surgery in the Netherlands (the HFO trial): a randomised, single-blind, adaptive non-inferiority trial. Lancet Neurol. 2022;21:982–993.

53. Wang S, Wang IZ, Bulacio JC, et al. Ripple classification helps to localize the seizure-onset zone in neocortical epilepsy. Epilepsia. 2013;54:370–376.

54. Gerstl JVE, Kiseleva A, Imbach L, Sarnthein J, Fedele T. High frequency oscillations in relation to interictal spikes in predicting postsurgical seizure freedom. Sci Rep. 2023;13:21313.

55. Shi W, Shaw D, Walsh KG, et al. Spike ripples localize the epileptogenic zone best: an international intracranial study. Brain. 2024;147:2496–2506.

56. Hufnagel A, Dümpelmann M, Zentner J, Schijns O, Elger CE. Clinical relevance of quantified intracranial interictal spike activity in presurgical evaluation of epilepsy. Epilepsia. 2000;41:467–478.

57. Nonoda Y, Miyakoshi M, Ojeda A, et al. Interictal high-frequency oscillations generated by seizure onset and eloquent areas may be differentially coupled with different slow waves. Clin Neurophysiol. 2016;127:2489–2499.

58. Iimura Y, Jones K, Takada L, et al. Strong coupling between slow oscillations and wide fast ripples in children with epileptic spasms: investigation of modulation index and occurrence rate. Epilepsia. 2018;59:544–554.

59. Motoi H, Miyakoshi M, Abel TJ, et al. Phase-amplitude coupling between interictal high-frequency activity and slow waves in epilepsy surgery. Epilepsia. 2018;59:1954–1965.

60. Roehri N, Pizzo F, Lagarde S, et al. High-frequency oscillations are not better biomarkers of epileptogenic tfpage than spikes. Ann Neurol. 2018;83:84–97.

61. Kural MA, Duez L, Sejer Hansen V, et al. Criteria for defining interictal epileptiform discharges in EEG: a clinical validation study. Neurology. 2020;94:e2139–e2147.

62. Kuroda N, Sonoda M, Miyakoshi M, et al. Objective interictal electrophysiology biomarkers optimize prediction of epilepsy surgery outcome. Brain Commun. 2021;3:fcab042.

63. Ma H, Wang Z, Li C, Chen J, Wang Y. Phase-amplitude coupling and epileptogenic zone localization of frontal epilepsy based on intracranial EEG. Front Neurol. 2021;12:718683.

64. Uda T, Kuki I, Inoue T, et al. Phase-amplitude coupling of interictal fast activities modulated by slow waves on scalp EEG and its correlation with seizure outcomes of disconnection surgery in children with intractable nonlesional epileptic spasms. J Neurosurg Pediatr. 2021;27:572–580.

65. Ricci L, Tamilia E, Mercier M, et al. Phase-amplitude coupling between low- and high-frequency activities as preoperative biomarker of focal cortical dysplasia subtypes. Clin Neurophysiol. 2023;150:40–48.

66. Sakakura K, Kuroda N, Sonoda M, et al. Developmental atlas of phase-amplitude coupling between physiologic high-frequency oscillations and slow waves. Nat Commun. 2023;14:6435.

67. Canolty RT, Edwards E, Dalal SS, et al. High gamma power is phase-locked to theta oscillations in human neocortex. Science. 2006;313:1626–1628.

68. Wada K, Sonoda M, Firestone E, et al. Sevoflurane-based enhancement of phase-amplitude coupling and localization of the epileptogenic zone. Clin Neurophysiol. 2022;134:1–8.

69. Firestone E, Sonoda M, Kuroda N, et al. Sevoflurane-induced high-frequency oscillations, effective connectivity and intraoperative classification of epileptic brain areas. Clin Neurophysiol. 2023;150:17–30.

70. Schreiber T. Measuring information transfer. Phys Rev Lett. 2000;85:461–464.

71. Ito S, Hansen ME, Heiland R, Lumsdaine A, Litke AM, Beggs JM. Extending transfer entropy improves identification of effective connectivity in a spiking cortical network model. PLoS One. 2011;6:e27431.

72. Uda H, Kuroda N, Firestone E, et al. Normative atlases of high-frequency oscillation and spike rates under sevoflurane anesthesia. Clin Neurophysiol. 2024;167:117–130.

73. Frauscher B, von Ellenrieder N, Zelmann R, et al. High-frequency oscialltions in the normal human brain. Ann Neurol. 2018;84:374–385.

74. Taylor PN, Papasavvas CA, Owen TW, et al. Normative brain mapping of interictal intracranial EEG to localize epileptogenic tfpage. Brain. 2022;145:939–949.

75. Malan TP Jr, DiNardo JA, Isner RJ, et al. Cardiovascular effects of sevoflurane compared with those of isoflurane in volunteers. Anesthesiology. 1995;83:918–928.

76. Nakai Y, Jeong JW, Brown EC, et al. Three- and four-dimensional mapping of speech and language in patients with epilepsy. Brain. 2017;140:1351–1370.

77. Pieters TA, Conner CR, Tandon N. Recursive grid partitioning on a cortical surface model: an optimized technique for the localization of implanted subdural electrodes. J Neurosurg. 2013;118:1086–1097.

78. Stolk A, Griffin S, van der Meij R, et al. Integrated analysis of anatomical and electrophysiological human intracranial data. Nat Protoc. 2018;13:1699–1723.

79. Miyakoshi M, Delorme A, Mullen T, Kojima K, Makeig S, Asano E. Automated detection of cross-frequency coupling in the electrocorticogram for clinical inspection. Annu Int Conf IEEE Eng Med Biol Soc. 2013;2013:3282–3285.

80. Oostenveld R, Fries P, Maris E, Schoffelen JM. FieldTrip: open source software for advanced analysis of MEG, EEG, and invasive electrophysiological data. Comput Intell Neurosci. 2011;2011:156869.

81. Delorme A, Makeig S. EEGLAB: an open source toolbox for analysis of singe-trial EEG dynamics including independent component analysis. J Neurosci Methods. 2004;134:9–21.

82. Desikan RS, Ségonne F, Fischl B, et al. An automated labeling system for subdividing the human cerebral cortex on MRI scans into gyral based regions of interest. Neuroimage. 2006;31:968–980.

83. Ghosh SS, Kakunoori S, Augustinack J, et al. Evaluating the validity of volume-based and surface-based brain image registration for developmental cognitive neuroscience studies in children 4 to 11 years of age. Neuroimage. 2010;53:85–93.

84. Silverstein BH, Asano E, Sugiura A, Sonoda M, Lee MH, Jeong JW. Dynamic tractography: integrating cortico-cortical evoked potentials and diffusion imaging. Neuroimage. 2020;215:116763.

85. Mitsuhashi T, Sonoda M, Sakakura K, et al. Dynamic tractography-based localization of spike sources and animation of spike propagations. Epilepsia. 2021;62:2372–2384.

86. Sonoda M, Silverstein BH, Jeong JW, et al. Six-dimensional dynamic tractography atlas of language connectivity in the developing brain. Brain. 2021;144:3340–3354.

87. Kitazawa Y, Sonoda M, Sakakura K, et al. Intra- and inter-hemispheric network dynamics supporting object recognition and speech production. Neuroimage. 2023;270:119954.

88. Hagmann P, Cammoun L, Gigandet X, et al. Mapping the structural core of human cerebral cortex. PLoS Biol. 2008;6:e159.

89. Yeh FC, Panesar S, Fernandes D, et al. Population-averaged atlas of the macroscale human structural connectome and its network topology. Neuroimage. 2018;178:57–68.

90. Harmony T. The functional significance of delta oscillations in cognitive processing. Front Integr Neurosci. 2013;7:83.

91. Jiruska P, Alvarado-Rojas C, Schevon CA, et al. Update on the mechanisms and roles of high-frequency oscillations in seizures and epileptic disorders. Epilepsia. 2017;58:1330–1339.

92. Tartaglia EM, Brunel N. Bistability and up/down state alternations in inhibition-dominated randomly connected networks of LIF neurons. Sci Rep. 2017;7:11916.

93. Jefferys JGR, Menendez de la Prida L, Wendling F, et al. Mechanisms of physiological and epileptic HFO generation. Prog Neurobiol. 2012;98:250–264.

94. Weiss SA, Fried I, Engel J Jr, et al. Fast ripples reflect increased excitability that primes epileptiform spikes. Brain Commun. 2023;5:fcad242.

95. Nickalls RWD, Mapleson WW. Age-related iso-MAC charts for isoflurane, sevoflurane and desflurane in man. Br J Anaesth. 2003;91:170–174.

96. Papadelis C, Tamilia E, Stufflebeam S, et al. Interictal High Frequency Oscillations Detected with Simultaneous Magnetoencephalography and Electroencephalography as Biomarker of Pediatric Epilepsy. J. Vis. Exp.2016;118:e54883.

97. Tamilia E, Dirodi M, Alhilani M, et al. Scalp ripples as prognostic biomarkers of epileptogenicity in pediatric surgery. Ann Clin Transl Neurol. 2020;7(3):329–342.

98. Dimakopoulos V, Gotman J, Klimes P, et al. Multicentre analysis of seizure outcome predicted by removal of high-frequency oscillations. Brain. 2024:awae361.

99. Weiss SA, Sheybani L, Seenarine N, et al. Delta oscillation coupled propagating fast ripples precede epileptiform discharges in patients with focal epilepsy. Neurobiol Dis. 2022;175:105928.

100. Tamrakar S, Iimura Y, Suzuki H, et al. Higher phase-amplitude coupling between ripple and slow oscillations indicates the distribution of epileptogenicity in temporal lobe epilepsy with hippocampal sclerosis. Seizure. 2022;100:1–7.

101. Dahal R, Tamura K, Pan D-S, et al. Effect of sevoflurane anesthesia on intraoperative spikes, high-frequency oscillations, and phase-amplitude coupling in MRI-normal hippocampus. J Clin Neurophysiol. 2024;41(7):589–596.

102. Axmacher N, Elger CE, Fell J. Ripples in the medial temporal lobe are relevant for human memory consolidation. Brain 2008;131(Pt 7):1806–1817.

103. Buzsáki G, Horváth Z, Urioste R, Hetke J, Wise K. High-frequency network oscillation in the hippocampus. Science 1992;256(5059):1025-1027. doi: 10.1126/science.1589772.

104. Buzsáki G. Hippocampal sharp wave-ripple: a cognitive biomarker for episodic memory and planning. Hippocampus 2015;25(10):1073–1188. doi: 10.1002/hipo.22488.

105. Engel AK, König P, Gray CM, Singer W. Stimulus-dependent neuronal oscillations in cat visual cortex: inter-columnar interaction as determined by cross-correlation analysis. Eur J Neurosci 1990;2(7):588–606.

106. Gray CM, König P, Engel AK, Singer W. Oscillatory responses in cat visual cortex exhibit inter-columnar synchronization which reflects global stimulus properties. Nature 1989;338(6213):334-337

107. Corona L, Tamilia E, Perry MS, et al. Non-invasive mapping of epileptogenic networks predicts surgical outcomes. Brain. 2023;146(5):1916–1931.

108. Rijal S, Corona L, Perry MS, et al. Functional connectivity discriminates epileptogenic states and predicts surgical outcome in children with drug resistant epilepsy. Sci Rep. 2023;13(1):9622.

109. Ntolkeras G, Makaram N, Bernabei M, et al. Interictal EEG source connectivity to localize the epileptogenic zone in patients with drug-resistant epilepsy: a machine learning approach. Epilepsia. 2024;65(4):944–960.

110. Lin J, Smith GC, Gliske SV, Zochowski M, Shedden K, Stacey WC. High-frequency oscillation network dynamics predict outcome in non-palliative epilepsy surgery. Brain Commun. 2024;6(1):fcae032.

111. Otárula KAG, von Ellenrieder N, Cuello-Oderiz C, Dubeau F, Gotman J. High-frequency oscillation networks and surgical outcome in adult focal epilepsy. Ann Neurol. 2019;85(4):485–494.

112. Yin C, Zhang X, Xiang J, et al. Altered effective connectivity network in patients with insular epilepsy: a high-frequency oscillations magnetoencephalography study. Clin Neurophysiol. 2020;131:377–384.

113. Emerson RG, Turner CA, Pedley TA, Walczak TS, Forgione M. Propagation patterns of temporal spikes. Electroencephalgr Clin Neurophysiol. 1995;94(5):338–348.

114. Alarcon G, Guy CN, Binnie CD, Walker SR, Elwes RD, Polkey CE. Intracerebral propagation of interictal activity in partial epilepsy: implications for source localization. J Neurol Neurosurg Psychiatry. 1994;57(4):435–449.

115. Matarrese MAG, Loppini A, Fabbri L, et al. Spike propagation mapping reveals effective connectivity and predicts surgical outcome in epilepsy. Brain. 2023;146:3898–3912.

116. Tamilia E, Matarrese MAG, Ntolkeras G, et al. Noninvasive mapping of ripple onset predicts outcome in epilepsy surgery. Ann Neurol. 2021;89:911–925.

117. Tamilia E, Park E-H, Percivati S, et al. Surgical resection of ripple onset predicts outcome in pediatric epilepsy. Ann Neurol. 2018;84:331–346.

118. Jahromi S, Matarrese MAG, Fabbri L, et al. Overlap of spike and ripple propagation onset predicts surgical outcome in epilepsy. Ann Clin Transl Neurol. 2024;11(10):2530–2547.

119. Shamas M, Yeh HJ, Fried I, Engel Jr J, Staba RJ. High-rate leading spikes in propagating spike sequences predict seizure outcome in surgical patients with temporal lobe epilepsy. Brain Commun. 2023;5(6):fcad289.

120. O’Hara NB, Lee MH, Juhász C, Asano E, Jeong JW. Diffusion tractography predicts propagated high-frequency activity during epileptic spasms. Epilepsia. 2022;63:1787–1798.

121. Azeem A, Abdallah C, von Ellenrieder N, Kosseifi CE, Frauscher B, Gotman J. Explaining slow seizure propagation with white matter tractography. Brain. 2024;147:3458–3470.

122. Azeem A, von Ellenrieder N, Royer J, Frauscher B, Bernhardt B, Gotman J. Integration of white matter architecture to stereo-EEG better describes epileptic spike propagation. Clin Neurophysiol. 2023;146:135–146.

123. Withers CP, Diamond JM, Yang B, et al. Identifying sources of human interictal discharges with travelling wave and white matter propagation. Brain. 2023;146:5168–5181.

124. Alkire MT, Hudetz AG, Tononi G. Consciousness and anesthesia. Science. 2008;322:876–880.

125. Ogawa SK, Tanaka E, Shin MC, Kotani N, Akaike N. Volatile anesthetic effects on isolated GABA synapses and extrasynaptic receptors. Neuropharmacology. 2011;60:701–710.

126. Xu W, Wang L, Yuan XS, et al. Sevoflurane depresses neurons in the medial parabrachial nucleus by potentiating postsynaptic GABA_A_ receptors and background potassium channels. Neuropharmacology. 2020;181:108249.

127. Mapelli J, Gandolfi D, Giuliani E, et al. The effects of the general anesthetic sevoflurane on neurotransmission: an experimental and computational study. Sci Rep. 2021;11:4335.

128. Llinás R, Jahnsen H. Electrophysiology of mammalian thalamic neurones in vitro. Nature. 1982;297:406–408.

129. Llinás R, Urbano FJ, Leznik E, Ramírez RR, van Marle HJF. Rhythmic and dysrhythmic thalamocortical dynamics: GABA systems and the edge effect. Trends Neurosci. 2005;28:325–333.

130. Crunelli V, Cope DW, Hughes SW. Thalamic T-type Ca2+ channels and NREM sleep. Cell Calcium. 2006;40:175–190.

131. Llinás R, Steriade M. Bursting of thalamic neurons and states of vigilance. J Neurophysiol. 2006;95:3297–3308.

132. Cruikshank SJ, Urabe H, Nurmikko AV, Connors BW. Pathway-specific feedforward circuits between thalamus and neocortex revealed by selective optical stimulation of axons. Neuron. 2010;65:230–245.

133. Crandall SR, Cruikshank SJ, Connors BW. A corticothalamic switch: controlling the thalamus with dynamic synapses. Neuron. 2015;86:768–782.

134. Suzuki SS, Smith GK. Spontaneous EEG spikes in the normal hippocampus. V. Effects of ether, urethane, pentobarbital, atropine, diazepam and bicuculline. Electroencephalogr Clin Neurophysiol. 1988;70:84–95.

135. Hapfelmeier G, Schneck H, Kochs E. Sevoflurane potentiates and blocks GABA-induced currents through recombinant alpha1beta2gamma2 GABAA receptors: implications for an enhanced GABAergic transmission. Eur J Anaesthesiol. 2001;18:377–383.

136. Eckle VS, Hauser S, Drexler B, Antkowiak B, Grasshoff C. Opposing actions of sevoflurane on GABAergic and glycinergic synaptic inhibition in the spinal ventral horn. PLoS One. 2013;8:e60286.

137. Calcagnotto ME, Paredes MF, Tihan T, Barbaro NM, Baraban SC. Dysfunction of synaptic inhibition in epilepsy associated with focal cortical dysplasia. J Neurosci. 2005;25:9649–9657.

138. Shao LR, Habela CW, Stafstrom CE. Pediatric epilepsy mechanisms: expanding the paradigm of excitation/inhibition imbalance. Children (Basel). 2019;6:23.

139. Schwartzkroin PA, Prince DA. Cellular and field potential properties of epileptogenic hippocampal slices. Brain Res. 1978;147:117–130.

140. Mercier MR, Dubarry A-S, Tadel F, et al. Advances in human intracranial electroencephalography research, guidelines, and good practices. NeuroImage. 2022;260:119438.

