## Supplementary figures for "Normative high-frequency oscillation phase-amplitude coupling and effective connectivity under sevoflurane"

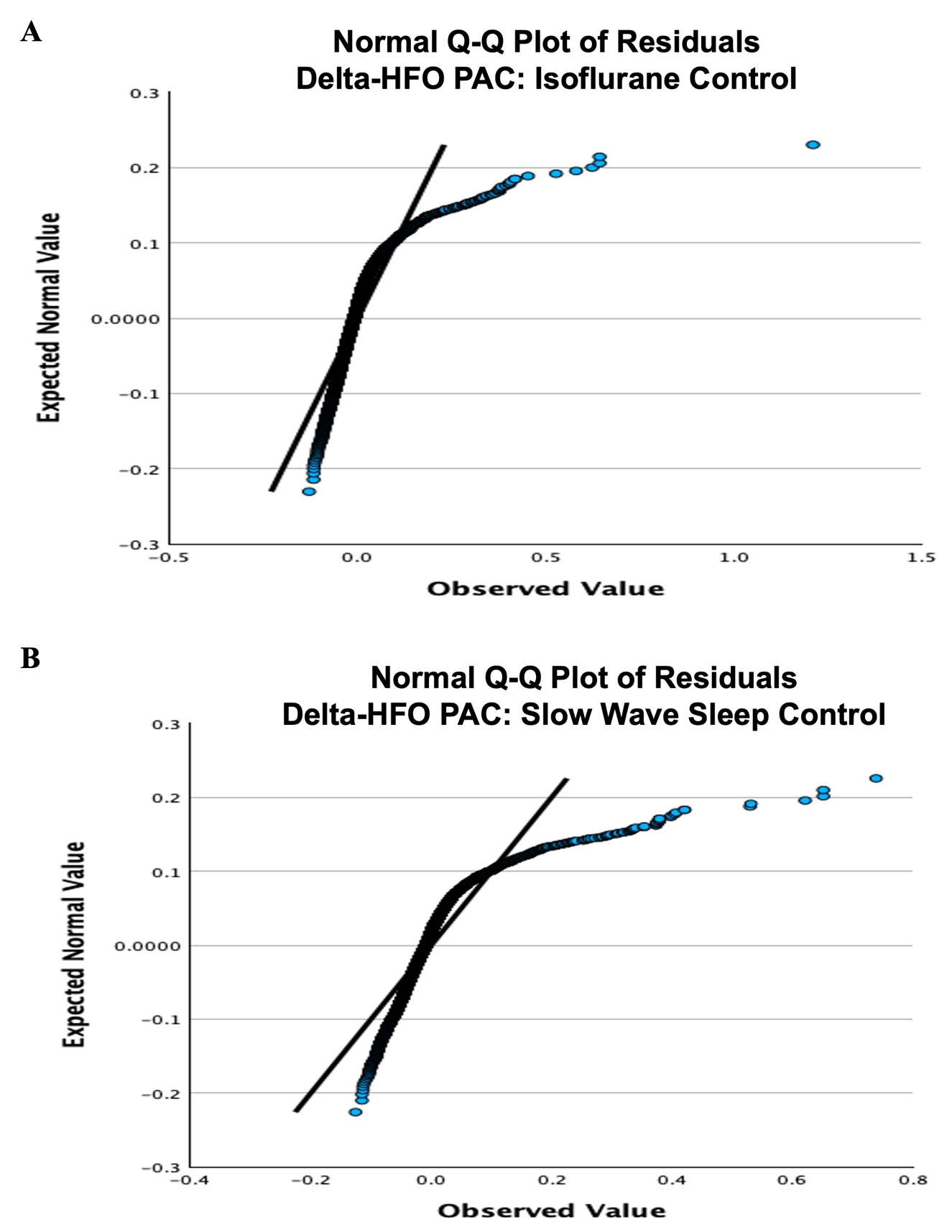

**Supplementary Figure 1. Q-Q plot of residuals for 3-4 Hz delta and HFO (80-300 Hz) phase-amplitude coupling response to sevoflurane. (A)** Q-Q plot of residuals for linear mixed model analysis of delta-HFO PAC, rated by modulation index, as a function of sevoflurane stage. Data from one-stage and two-stage patients was used. The isoflurane stage was the reference period for the linear mixed model. **(B)** Q-Q plot of residuals for linear mixed model analysis of delta-HFO PAC, rated by modulation index, as a function of sevoflurane stage. Data from two-stage patients was used. The slow wave sleep stage was the reference period for the linear mixed model. HFO = high-frequency oscillation; PAC = phase-amplitude coupling; Q-Q = quantile-quantile.

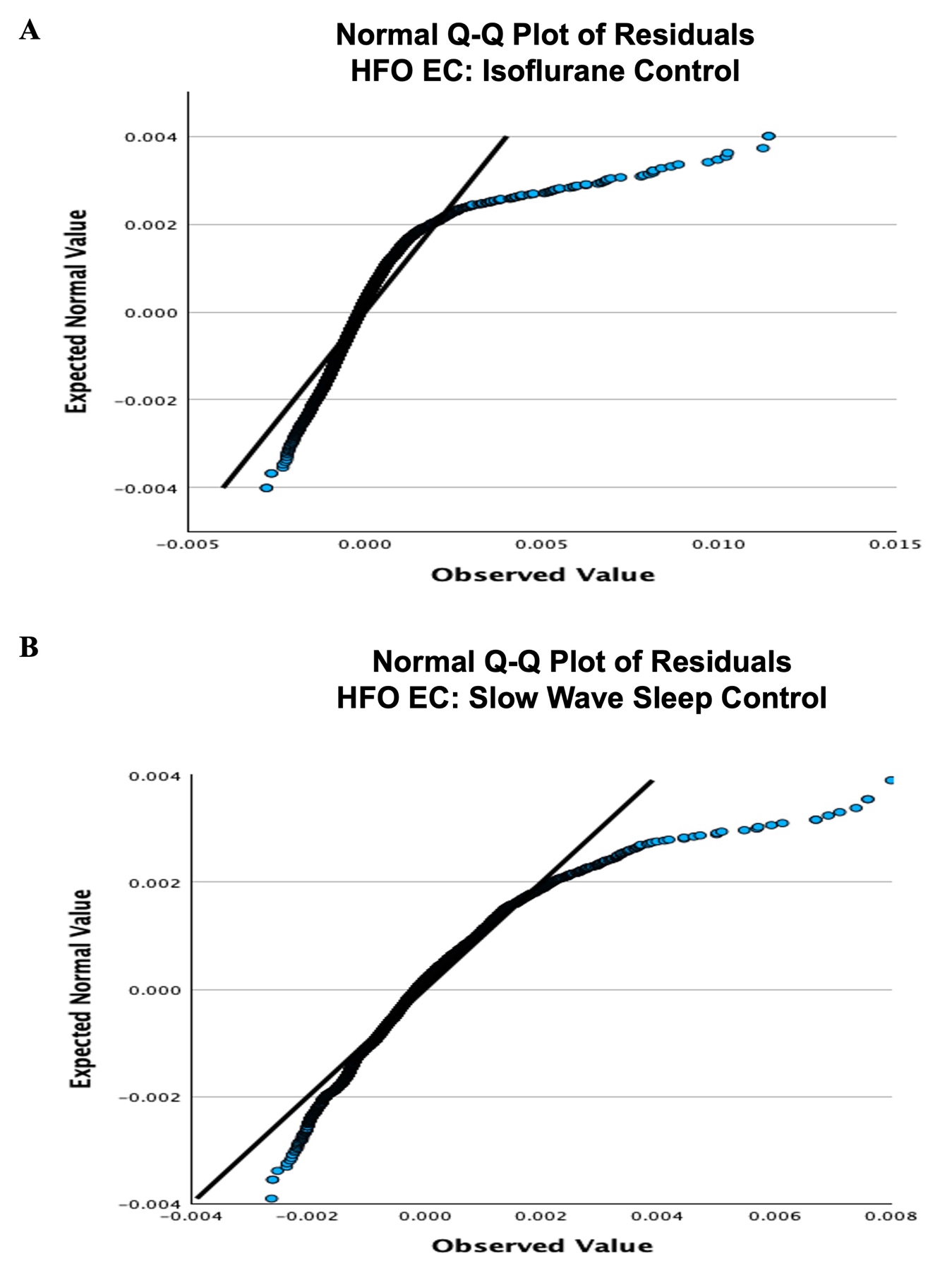

**Supplementary Figure 2. Q-Q plot of residuals for HFO (80-300 Hz) effective connectivity response to sevoflurane. (A)** Q-Q plot of residuals for linear mixed model analysis of HFO EC, rated by transfer entropy, as a function of sevoflurane stage. HFO EC values from one-stage and two-stage patients were derived from a spectral amplitude time series z-scored against the mean and standard deviation of the isoflurane control. In addition, the isoflurane stage was the reference period for the linear mixed model. **(B)** Q-Q plot of residuals for linear mixed model analysis of HFO EC, rated by transfer entropy, as a function of sevoflurane stage. HFO EC values from two-stage patients were derived from a spectral amplitude time series z-scored against the mean and standard deviation of the slow wave sleep control. In addition, the slow wave sleep stage was the reference period for the linear mixed model. EC = effective connectivity; HFO = high-frequency oscillation; Q-Q = quantile-quantile.

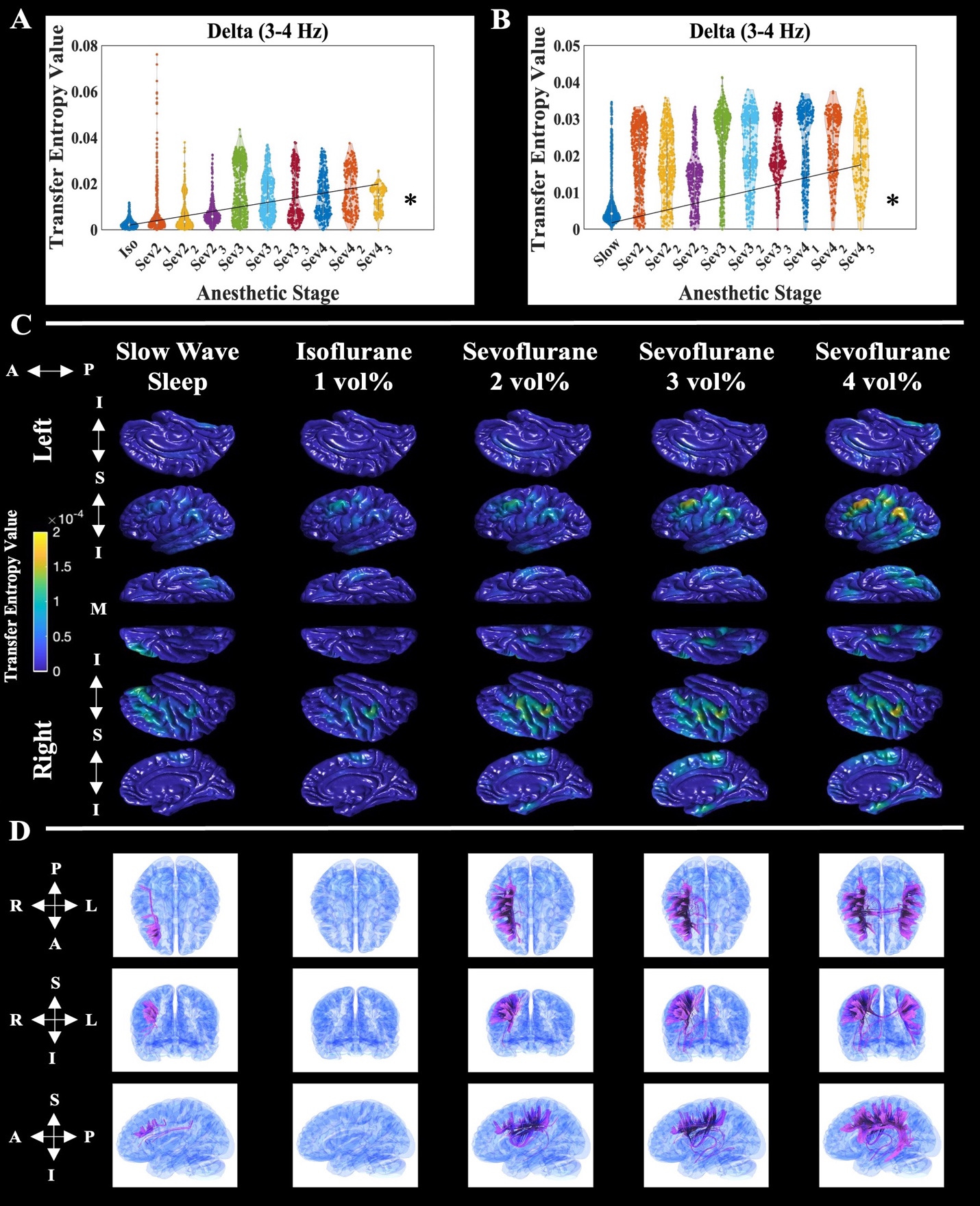

**Supplementary Figure 3. Normative sevoflurane distribution of delta (3-4 Hz) effective connectivity.** Mathematical distribution of ECoG/iEEG-defined delta effective connectivity – rated by transfer entropy - from all pooled normative electrode sites, at each anesthetic stage. The left panel **(A)** shows delta EC data from one-stage and two-stage patients derived from a spectral amplitude time series z-scored in relation to the mean and standard deviation of the isoflurane control (iso; *n = 963* normative electrode sites) and a linear mixed model that uses the isoflurane period as the reference. The right panel **(B)** shows delta EC data from two-stage patients derived from a spectral amplitude time series z-scored in relation to the mean and standard deviation of the slow wave sleep control (slow; *n = 816* normative electrode sites) and a linear mixed model that uses the slow wave sleep period as the reference. In addition, ‘Sev2_1’ denotes the first minute of sevoflurane at a concentration of two-volume-percent, and so on. The black trend line anchored at the origin depicts the relationship between delta effective connectivity and anesthetic stage: determined by linear mixed model analysis. The asterisks represent a significant effect (Bonferroni corrected p < 0.05) of anesthetic stage on biomarker value, via linear mixed model analysis. **(C)** Anatomical distribution of normative delta EC interpolated onto the FreeSurfer average cortical surface. The data represents normative electrode sites pooled from the eight patients who underwent both extraoperative iEEG recording during slow wave sleep, as well as intraoperative ECoG at every anesthetic stage (*n = 431* normative electrode sites). Each column represents a different anesthetic stage. The left and right hemispheres are grouped in the first and last three rows, respectively. Within each hemisphere group, single rows represent different cortical surfaces. For the left hemisphere: top row-sagittal-medial, middle row-sagittal-lateral, and bottom row-axial-inferior. The orientation row order is opposite for the right hemisphere. The anterior – posterior (‘A 🡨🡪 P’) orientation is listed for all views. In addition, the inferior and superior orientations for the sagittal images are depicted (‘I 🡨🡪S 🡨🡪 I’), as well as the medial (‘M’) marker for the inferior-axial view. Hotter colors represent sites with relatively higher delta EC values and vice versa. The spatial coverage of intracranial electrodes is presented in Figure 2. In areas without intracranial electrode coverage, the absence of biomarker values should be interpreted as a lack of intracranial EEG signal sampling. **(D)** Dynamic tractography depicts diffusion weighted-imaging (DWI) white matter streamlines connecting cortical sites with significantly elevated ECoG/iEEG-defined delta EC (i.e., greater than three standard deviations above the slow wave sleep mean). Each column represents a different anesthetic stage, and each row, from top to bottom, shows a different brain view: axial, coronal, and sagittal, respectively. The orientation for each view is included to the left of the corresponding row. A = anterior; DWI = diffusion-weighted imaging; EC = effective connectivity; extraoperative iEEG = extraoperative intracranial electroencephalography; I = inferior; intraoperative ECoG = intraoperative electrocorticography; L = left; M = medial; P = posterior; R = right; S = superior.

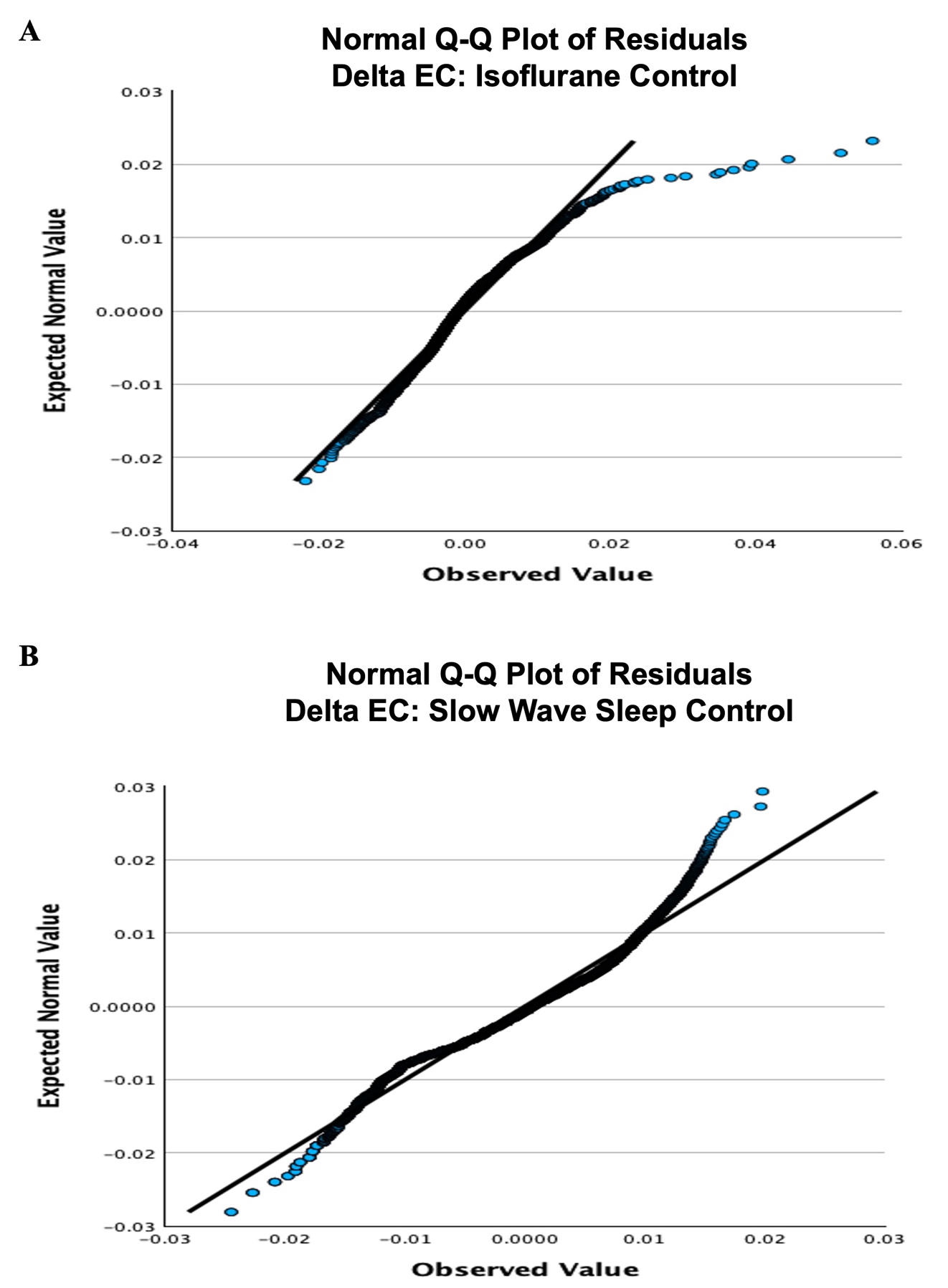

**Supplementary Figure 4. Q-Q plot of residuals for delta (3-4 Hz) effective connectivity response to sevoflurane. (A)** Q-Q plot of residuals for linear mixed model analysis of delta EC, rated by transfer entropy, as a function of sevoflurane stage. Delta EC values from one-stage and two-stage patients were derived from a spectral amplitude time series z-scored against the mean and standard deviation of the isoflurane control. In addition, the isoflurane stage was the reference period for the linear mixed model. **(B)** Q-Q plot of residuals for linear mixed model analysis of delta EC, rated by transfer entropy, as a function of sevoflurane stage. Delta EC values from two-stage patients were derived from a spectral amplitude time series z-scored against the mean and standard deviation of the slow wave sleep control. In addition, the slow wave sleep stage was the reference period for the linear mixed model. Q-Q = quantile-quantile.

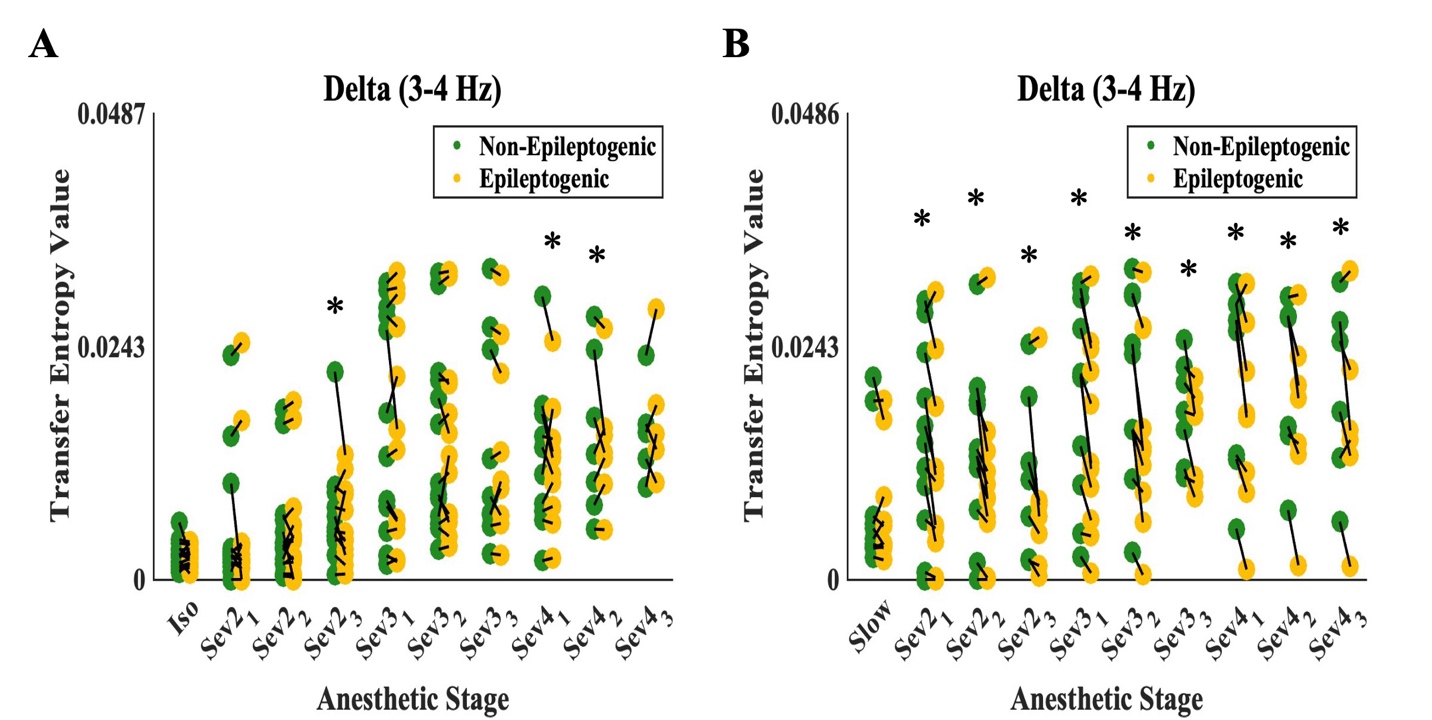

**Supplementary Figure 5. Epileptogenic characterization using sevoflurane-activated delta (3-4 Hz) effective connectivity.** Delta effective connectivity (rated by transfer entropy) values derived from a spectral amplitude time series z-scored in relation to the mean and standard deviation of the **(A)** isoflurane (*n = 1,608* total electrode sites from one-stage and two-stage patients) and **(B)** slow wave sleep (*n = 1,344* total electrode sites from two-stage patients) controls, at each anesthetic stage. For a given patient, the green and yellow dots represent the average value of all non-epileptogenic and epileptogenic sites, respectively. In each anesthetic stage, the black lines connect a pair of green-yellow dots from the same patient. The asterisks denote binary logistic mixed model significance (Bonferroni corrected p < 0.05) for characterizing epileptogenicity via delta effective connectivity levels. On the x-axis, ‘Sev2_1’ denotes the first minute of sevoflurane at a concentration of two-volume-percent, and so on. ‘Iso’ refers to the isoflurane period, while ‘Slow’ refers to the slow-wave sleep period.

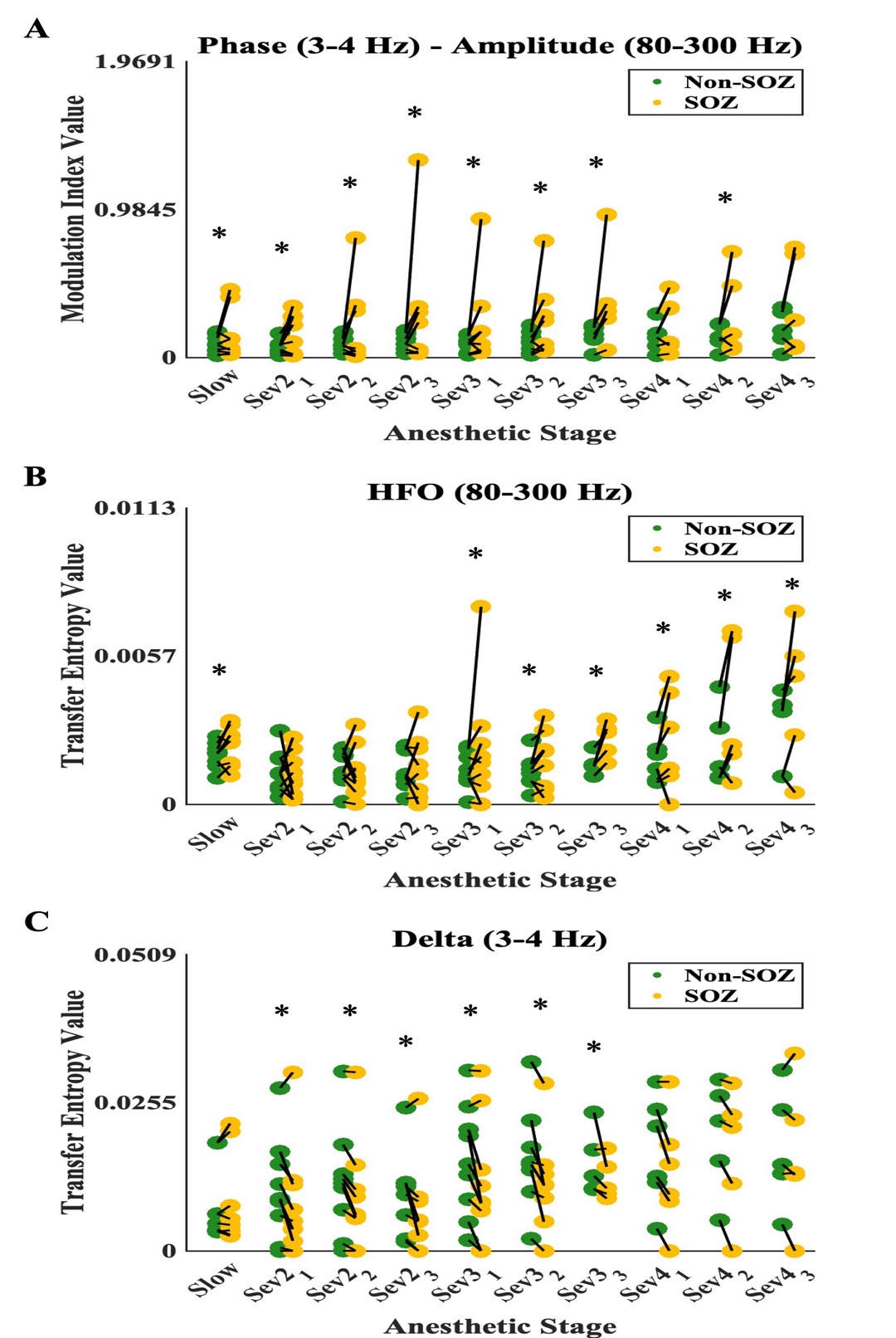

**Supplementary Figure 6. Seizure onset zone characterization via ECoG/iEEG biomarkers.** ECoG/iEEG-based biomarker values from two-stage patients (*n = 1,344* total electrode sites), for **(A)** delta-HFO phase-amplitude coupling (rated by modulation index), **(B)** HFO effective connectivity (rated by transfer entropy), and **(C)** delta effective connectivity (rated by transfer entropy), at each anesthetic stage. Effective connectivity values were derived from a spectral amplitude time series z-scored based on the mean and standard deviation of the slow wave sleep control. For a given patient, the green and yellow dots represent the average value of all non-seizure onset zone (non-SOZ) and seizure onset zone (SOZ) sites, respectively. In each anesthetic stage, the black lines connect a pair of green-yellow dots from the same patient. The asterisks denote binary logistic mixed model significance (Bonferroni corrected p < 0.05) for characterizing SOZ status via ECoG/iEEG biomarker levels. On the x-axis, ‘Sev2_1’ denotes the first minute of sevoflurane at a concentration of two-volume-percent, and so on. ‘Slow’ represents the slow wave sleep period. Extraoperative iEEG = extraoperative intracranial electroencephalography; HFO = high-frequency oscillation; intraoperative ECoG = intraoperative electrocorticography; SOZ = seizure onset zone.
