## Supplementary tables for "Normative high-frequency oscillation phase-amplitude coupling and effective connectivity under sevoflurane"

**Supplementary Table 1. Children’s Hospital of Michigan recurrent seizure outcome patient profiles.**

| **Patient Number** | **Age (years)** | **Biological Sex** | **Number Anti-Seizure Medications** | **Number Electrodes** | **One- versus Two-Stage Surgery** | **MRI Finding** |
| --- | --- | --- | --- | --- | --- | --- |
| s1 | 20 | M | 2 | 118 | 2 | Nonlesional |
| s2 | 8 | F | 3 | 126 | 2 | Dysplasia |
| s3 | 6 | M | 3 | 134 | 2 | Focal atrophy |
| s4 | 6 | M | 3 | 134 | 2 | Dysplasia |

**Supplementary Table 2. Normative sites: linear mixed model analysis of the high-frequency oscillation effective connectivity (80 – 300 Hz) response to anesthesia.**

| **Biomarker** | **Control vs. Exposure** | **t-value** | **Uncorrected p-value** | **Bonferroni Corrected p-value** | **Fixed-Effect Estimate** |
| --- | --- | --- | --- | --- | --- |
| *HFO aEC:*  *80-300 Hz*  (df = 1,239) | slow vs. iso | 1.314 | 0.189 | > 0.99 | 9.132E-5 |
| ***HFO aEC:***  ***80-300 Hz***  **(df = 4,008)** | **iso vs. sev** | **7.688** | **1.86E-14** | **1.12E-13** | **5.850E-5** |
| ***HFO aEC:***  ***80-300 Hz***  **(df = 4,419)** | **slow vs. sev** | **6.408** | **1.63E-10** | **9.78E-10** | **3.966E-5** |
| *HFO eEC:*  *80-300 Hz*  (df = 1,239) | slow vs. iso | 1.016 | 0.310 | > 0.99 | 7.255E-5 |
| ***HFO eEC:***  ***80-300 Hz***  **(df = 4,008)** | **iso vs. sev** | **8.237** | **2.37E-16** | **1.42E-15** | **6.308E-5** |
| ***HFO eEC:***  ***80-300 Hz***  **(df = 4,419)** | **slow vs. sev** | **6.864** | **7.62E-12** | **4.57E-11** | **4.250E-5** |

Effects of anesthetic stage (significance at Bonferroni corrected p < 0.05 for 6 comparisons, as in the main analysis) on HFO effective connectivity (rated by transfer entropy). Significant values are bolded and quantified in the table. Afferent EC (‘aEC’) refers to connections converging on a site, while efferent EC (‘eEC’) refers to connections emanating from a site. In the *Control vs. Exposure* column, the first stage refers to the reference condition that all spectral amplitude values were z-scored against. It also denotes the reference condition used in linear mixed model analysis. For example, ‘iso vs. sev’ means that all spectral amplitude values used to calculate transfer entropy were z-scored against the mean and standard deviation of the isoflurane control, and the linear mixed model was assessed at all sevoflurane stages, in relation to the isoflurane control.

aEC = afferent effective connectivity; df = degrees of freedom; eEC = efferent effective connectivity; HFO = high-frequency oscillation; iso = isoflurane; sev = sevoflurane; slow = slow wave sleep.

**Supplementary Table 3. Normative sites: linear mixed model analysis of the delta oscillation effective connectivity (3 – 4 Hz) response to anesthesia.**

| **Biomarker** | **Control vs. Exposure** | **t-value** | **Uncorrected p-value** | **Bonferroni Corrected p-value** | **Fixed-Effect Estimate** |
| --- | --- | --- | --- | --- | --- |
| ***Delta EC:***  ***3-4 Hz***  **(df = 1,239)** | **slow vs. iso** | **10.387** | **2.74E-24** | **1.64E-23** | **0.004** |
| ***Delta EC:***  ***3-4 Hz***  **(df =4,001)** | **iso vs. sev** | **48.800** | ***** | ***** | **0.0020** |
| ***Delta EC:***  ***3-4 Hz***  **(df = 4,417)** | **slow vs. sev** | **39.378** | **5.92E-291** | **3.55E-290** | **0.0018** |
| ***Delta aEC*:**  **3-4 Hz**  **(df = 1,239)** | **slow vs. iso** | **10.055** | **6.45E-23** | **3.87E-22** | **0.004** |
| ***Delta aEC:***  ***3-4 Hz***  **(df = 4,001)** | **iso vs. sev** | **46.750** | ***** | ***** | **0.0020** |
| ***Delta aEC:***  ***3-4 Hz***  **(df = 4,417)** | **slow vs. sev** | **37.450** | **7.97E-267** | **4.78E-266** | **0.0018** |
| ***Delta eEC:***  ***3-4 Hz***  **(df = 1,236)** | **slow vs. iso** | **10.268** | **8.55E-24** | **5.13E-22** | **0.004** |
| ***Delta eEC:***  ***3-4 Hz***  **(df = 4,001)** | **iso vs. sev** | **47.956** | ***** | ***** | **0.0020** |
| ***Delta eEC:***  ***3-4 Hz***  **(df = 4,416)** | **slow vs. sev** | **39.803** | **2.38E-296** | **1.43E-295** | **0.0018** |

Effects of anesthetic stage (significance at Bonferroni corrected p < 0.05 for 6 comparisons, as in the main analysis) on delta effective connectivity (rated by transfer entropy). Significant values are bolded and quantified in the table. Afferent EC (‘aEC’) refers to connections converging on a site, while efferent EC (‘eEC’) refers to connections emanating from a site. The ‘EC’ rows represent the average of both the ‘aEC’ and ‘eEC’. In the *Control vs. Exposure* column, the first stage refers to the reference condition that all spectral amplitude values were z-scored against. It also denotes the reference condition used in linear mixed model analysis. For example, ‘iso vs. sev’ means that all spectral amplitude values used to calculate transfer entropy were z-scored against the mean and standard deviation of the isoflurane control, and the linear mixed model was assessed at all sevoflurane stages, in relation to the isoflurane control.

*: SPSS only registered p-value as “ < 0.001” due to incredibly small value.

aEC = afferent effective connectivity; df = degrees of freedom; EC = effective connectivity; eEC = efferent effective connectivity; iso = isoflurane; sev = sevoflurane; slow = slow wave sleep.

**Supplementary Table 4. Binary logistic mixed model characterization of epileptogenicity based on the delta effective connectivity (3 - 4 Hz) response to anesthesia.**

| **Fixed effect variable*** | **Number of patients included in the analysis **** | **DF** | **t-value** | **Uncorrected p-value** | **Bonferroni-corrected**  **p-value***** | **OR** |
| --- | --- | --- | --- | --- | --- | --- |
| Delta EC during Isoflurane  (1 vol%) | 18: One-stage  and Two-stage | 994 | 0.476 | 0.634 | > 0.99 | 1.037E10 |
| Delta EC during Slow Wave Sleep | 11: Two-stage only | 1,279 | -1.355 | 0.176 | > 0.99 | 1.518E-9 |
| **Delta EC during the 1st minute of Sevoflurane**  **(2 vol%)** | 18: One-stage  and Two-stage | 858 | -0.695 | 0.487 | > 0.99 | 2.561E-7 |
|  | **11: Two-stage only** | **941** | **-8.197** | **8.88E-16** | **3.55E-14** | **7.595E-47** |
| **Delta EC during the 2nd minute of Sevoflurane**  **(2 vol%)** | 18: One-stage  and Two-stage | 854 | 1.363 | 0.173 | > 0.99 | 9.272E17 |
|  | **11: Two-stage only** | **928** | **-8.449** | **2.22E-16** | **8.88E-15** | **3.171E-49** |
| **Delta EC during the 3rd minute of Sevoflurane**  **(2 vol%)** | **18: One-stage**  **and Two-stage** | **652** | **-3.449** | **0.00060** | **0.024** | **1.076E-63** |
|  | **11: Two-stage only** | **643** | **-9.252** | ******** | ******** | **7.302E-108** |
| **Delta EC during the 1st minute of Sevoflurane**  **(3 vol%)** | 18: One-stage  and Two-stage | 815 | -1.737 | 0.083 | > 0.99 | 2.160E-14 |
|  | **11: Two-stage only** | **898** | **-10.610** | ******** | ******** | **1.595E-55** |
| **Delta EC during the 2nd minute of Sevoflurane**  **(3 vol%)** | 18: One-stage  and Two-stage | 744 | -1.641 | 0.101 | > 0.99 | 2.990E-18 |
|  | **11: Two-stage only** | **816** | **-9.287** | ******** | ******** | **1.204E-57** |
| **Delta EC during the 3rd minute of Sevoflurane**  **(3 vol%)** | 18: One-stage  and Two-stage | 449 | 0.609 | 0.543 | > 0.99 | 9.669E10 |
|  | **11: Two-stage only** | **547** | **-6.523** | **1.57E-10** | **6.28E-9** | **7.919E-61** |
| **Delta EC during the 1st minute of Sevoflurane**  **(4 vol%)** | **18: One-stage**  **and Two-stage** | **512** | **-3.935** | **9.0E-5** | **3.6E-3** | **2.387E-44** |
|  | **11: Two-stage only** | **604** | **-8.291** | **6.66E-16** | **2.66E-14** | **4.326E-72** |
| **Delta EC during the 2nd minute of Sevoflurane**  **(4 vol%)** | **18: One-stage**  **and Two-stage** | **411** | **-5.309** | **1.80E-7** | **7.2E-6** | **5.505E-66** |
|  | **11: Two-stage only** | **538** | **-7.099** | **4.00E-12** | **1.6E-10** | **1.340E-63** |
| **Delta EC during the 3rd minute of Sevoflurane**  **(4 vol%)** | 18: One-stage  and Two-stage | 315 | 2.666 | 0.008 | 0.32 | 2.682E44 |
|  | **11: Two-stage only** | **460** | **-5.688** | **2.29E-8** | **9.16E-7** | **7.992E-54** |

*: The delta effective connectivity values (rated by transfer entropy), as an ECoG/iEEG biomarker, was incorporated into the multivariate binary logistic mixed model analysis, which included eight fixed effect predictors: [1] age, [2] biological sex, [3] sampled hemisphere, [4] presence of MRI lesion, [5] presence of daily seizures, [6] number of anti-seizure medications, [7] one- versus two-stage surgery, and [8] biomarker value. Patient and intercept were considered random effects variables. The binary response variable was epileptogenic- or non-epileptogenic status of electrodes (1 or 0, respectively).

**: In the analysis including both one-stage and two-stage surgery patients, transfer entropy was calculated using a spectral amplitude time series that was normalized via z-scoring based on the mean and standard deviation of the isoflurane (1 vol%) period. In the analysis with only two-stage surgery patients, transfer entropy was calculated using a spectral amplitude time series that was normalized via z-scoring based on the mean and standard deviation of the slow wave sleep period.

***: Bonferroni correction was applied due to the 40 analyses conducted in the main analysis, and significant time points are bolded (Bonferroni corrected p-value < 0.05).

****: SPSS only registered p-value as “ < 0.001” due to incredibly small value.

DF: degrees of freedom; EC = effective connectivity; extraoperative iEEG = extraoperative intracranial electroencephalography; intraoperative ECoG = intraoperative electrocorticography; OR: odds ratio.

**Supplementary Table 5. Cohen’s *d** effect size of biomarker ability to characterize epileptogenicity in response to anesthesia.**

| **ECoG/iEEG biomarker** | **Number of patients included in the analysis **** | **Isoflurane (1 vol%)** | **Slow wave sleep** | **Sevoflurane (2 vol%)** | **Sevoflurane (3 vol%)** | **Sevoflurane (4 vol%)** |
| --- | --- | --- | --- | --- | --- | --- |
| Delta-HFO phase-amplitude coupling  (modulation index) | 18: One-stage  and Two-stage | 0.0106 | ** | 0.1381 | 0.2472 | 0.2328 |
|  | 11: Two-stage only | ** | 0.1619 | 0.2658 | 0.3665 | 0.3608 |
| HFO (80-300 Hz) effective connectivity  (transfer entropy) | 18: One-stage  and Two-stage | 0.1235 | ** | 0.1384 | 0.7250 | 0.6921 |
|  | 11: Two-stage only | ** | 0.2221 | 0.0609 | 0.3430 | 0.5616 |
| Delta (3-4 Hz) effective connectivity  (transfer entropy) | 18: One-stage  and Two-stage | 0.0470 | ** | 0.1316 | 0.0522 | 0.1934 |
|  | 11: Two-stage only | ** | 0.0270 | 0.6181 | 0.6709 | 0.6698 |

*: Cohen’s *d* was calculated to estimate the effect size of the influence of various anesthetic conditions on biomarker values between the epileptogenic and non-epileptogenic sites. For each patient and each one-minute segment of a given anesthetic stage, Cohen’s *d* was calculated as the absolute value of, [(the average biomarker value in epileptogenic sites) minus (the average biomarker value in non-epileptogenic sites)] divided by (the standard deviation across all sites). Values for all patients and all three one-minute segments, for a given anesthetic stage, were averaged to produce one effect size value for the corresponding anesthetic stage.

**: In the analysis including both one-stage and two-stage surgery patients, transfer entropy was calculated using a spectral amplitude time series that was normalized via z-scoring based on the mean and standard deviation of the isoflurane (1 vol%) period. In the analysis with only two-stage surgery patients, transfer entropy was calculated using a spectral amplitude time series that was normalized via z-scoring based on the mean and standard deviation of the slow wave sleep period. The same patient distributions were also used for the two different modulation index calculations. The control period for the one-stage plus two-stage patients was the isoflurane stage. The control period for the two-stage only patients was the slow wave sleep period.

Extraoperative iEEG = extraoperative intracranial electroencephalography; HFO = high-frequency oscillation; intraoperative ECoG = intraoperative electrocorticography.

**Supplementary Table 6. Cohen’s *d** effect size of biomarker ability to characterize seizure onset zone status in response to anesthesia.**

| **ECoG/iEEG biomarker** | **Number of patients included in the analysis **** | **Slow wave sleep** | **Sevoflurane (2 vol%)** | **Sevoflurane (3 vol%)** | **Sevoflurane (4 vol%)** |
| --- | --- | --- | --- | --- | --- |
| Delta-HFO phase-amplitude coupling  (modulation index) | 11: Two-stage only | 0.3820 | 0.4170 | 1.0144 | 0.6431 |
| HFO (80-300 Hz) effective connectivity  (transfer entropy) | 11: Two-stage only | 0.5216 | 0.0595 | 0.6314 | 0.7247 |
| Delta (3-4 Hz) effective connectivity  (transfer entropy) | 11: Two-stage only | 0.1539 | 0.5189 | 0.6320 | 0.4406 |

*: Cohen’s *d* was calculated to estimate the effect size of the influence of various anesthetic conditions on biomarker values between the SOZ sites and non-SOZ sites. For each patient and each one-minute segment of a given anesthetic stage, Cohen’s *d* was calculated as the absolute value of, [(the average biomarker value in SOZ sites) minus (the average biomarker value in non-SOZ sites)] divided by (the standard deviation across all sites). Values for all patients and all three one-minute segments, for a given anesthetic stage, were averaged to produce one effect size value for the corresponding anesthetic stage.

**: In the analysis with only two-stage surgery patients, transfer entropy was calculated using a spectral amplitude time series that was normalized via z-scoring based on the mean and standard deviation of the slow wave sleep period. The same patient distribution was also used for the modulation index calculation. The control period for the two-stage only patients was the slow wave sleep period.

Extraoperative iEEG = extraoperative intracranial electroencephalography; HFO = high-frequency oscillation; intraoperative ECoG = intraoperative electrocorticography; SOZ = seizure onset zone.

**Supplementary Table 7. Binary logistic mixed model characterization of seizure onset zone based on the delta (3-4 Hz) and high-frequency oscillation (80-300 Hz) phase-amplitude coupling response to anesthesia.**

| **Fixed effect variable*** | **Number of patients included in the analysis** | **DF** | **t-value** | **Uncorrected p-value** | **Bonferroni-corrected p-value**** | **OR** |
| --- | --- | --- | --- | --- | --- | --- |
| **Delta-HFO PAC during Slow Wave Sleep** | **11: Two-stage only** | **1,279** | **5.654** | **1.94E-8** | **7.76E-7** | **86.751** |
| **Delta-HFO PAC during the 1st minute of Sevoflurane**  **(2 vol%)** | **11: Two-stage only** | **959** | **6.201** | **8.34E-10** | **3.34E-8** | **782.293** |
| **Delta-HFO PAC during the 2nd minute of Sevoflurane**  **(2 vol%)** | **11: Two-stage only** | **951** | **5.402** | **8.35E-8** | **3.34E-6** | **26.436** |
| **Delta-HFO PAC during the 3rd minute of Sevoflurane**  **(2 vol%)** | **11: Two-stage only** | **656** | **5.279** | **1.77E-7** | **7.08E-6** | **25.498** |
| **Delta-HFO PAC during the 1st minute of Sevoflurane**  **(3 vol%)** | **11: Two-stage only** | **915** | **4.809** | **1.78E-6** | **7.12E-5** | **31.488** |
| **Delta-HFO PAC during the 2nd minute of Sevoflurane**  **(3 vol%)** | **11: Two-stage only** | **831** | **5.102** | **4.17E-7** | **1.67E-5** | **26.345** |
| **Delta-HFO PAC during the 3rd minute of Sevoflurane**  **(3 vol%)** | **11: Two-stage only** | **555** | **5.403** | **9.73E-8** | **3.89E-6** | **18.198** |
| Delta-HFO PAC during the 1st minute of Sevoflurane  (4 vol%) | 11: Two-stage only | 621 | 2.520 | 0.012 | 0.48 | 3.766 |
| **Delta-HFO PAC during the 2nd minute of Sevoflurane**  **(4 vol%)** | **11: Two-stage only** | **555** | **4.001** | **7.18E-5** | **0.0029** | **16.064** |
| Delta-HFO PAC during the 3rd minute of Sevoflurane  (4 vol%) | 11: Two-stage only | 472 | 3.054 | 0.002 | 0.08 | 3.610 |

*: The delta-HFO PAC value (rated by modulation index), as an ECoG/iEEG biomarker, was incorporated into the multivariate binary logistic mixed model analysis, which included eight fixed effect predictors: [1] age, [2] biological sex, [3] sampled hemisphere, [4] presence of MRI lesion, [5] presence of daily seizures, [6] number of anti-seizure medications, [7] one- versus two-stage surgery, and [8] biomarker value. Patient and intercept were considered random effects variables. The binary response variable was SOZ or non-SOZ status of electrodes (1 or 0, respectively).

**: Bonferroni correction was applied due to the 40 analyses conducted in the main analysis, and significant time points are bolded (Bonferroni corrected p-value < 0.05).

DF: degrees of freedom; extraoperative iEEG = extraoperative intracranial electroencephalography; HFO = high-frequency oscillation; intraoperative ECoG = intraoperative electrocorticography; OR: odds ratio; PAC = phase-amplitude coupling; SOZ: seizure onset zone.

**Supplementary Table 8. Binary logistic mixed model characterization of seizure onset zone based on the high-frequency oscillation effective connectivity (80 – 300 Hz) response to anesthesia.**

| **Fixed effect variable*** | **Number of patients included in the analysis **** | **DF** | **t-value** | **Uncorrected p-value** | **Bonferroni-corrected p-value***** | **OR** |
| --- | --- | --- | --- | --- | --- | --- |
| **HFO EC during Slow Wave Sleep** | **11: Two-stage only** | **1,279** | **4.502** | **7.35E-6** | **0.00029** | ******** |
| HFO EC during the 1st minute of Sevoflurane  (2 vol%) | 11: Two-stage only | 941 | 1.441 | 0.150 | > 0.99 | 3.559E91 |
| HFO EC during the 2nd minute of Sevoflurane  (2 vol%) | 11: Two-stage only | 928 | 1.874 | 0.061 | > 0.99 | 7.073E133 |
| HFO EC during the 3rd minute of Sevoflurane  (2 vol%) | 11: Two-stage only | 643 | 2.047 | 0.041 | > 0.99 | 1.819E155 |
| **HFO EC during the 1st minute of Sevoflurane**  **(3 vol%)** | **11: Two-stage only** | **898** | **6.858** | **1.30E-11** | **5.2E-10** | ******** |
| **HFO EC during the 2nd minute of Sevoflurane**  **(3 vol%)** | **11: Two-stage only** | **816** | **5.097** | **4.29E-7** | **1.72E-5** | ******** |
| **HFO EC during the 3rd minute of Sevoflurane**  **(3 vol%)** | **11: Two-stage only** | **547** | **5.744** | **1.54E-8** | **6.16E-7** | ******** |
| **HFO EC during the 1st minute of Sevoflurane**  **(4 vol%)** | **11: Two-stage only** | **604** | **4.661** | **3.88E-6** | **0.00016** | **9.397E210** |
| **HFO EC during the 2nd minute of Sevoflurane**  **(4 vol%)** | **11: Two-stage only** | **538** | **4.230** | **2.75E-5** | **0.0011** | **8.698E185** |
| **HFO EC during the 3rd minute of Sevoflurane**  **(4 vol%)** | **11: Two-stage only** | **460** | **4.090** | **5.09E-5** | **0.0020** | **1.381E167** |

*: The HFO effective connectivity value (rated by transfer entropy), as an ECoG/iEEG biomarker, was incorporated into the multivariate binary logistic mixed model analysis, which included eight fixed effect predictors: [1] age, [2] biological sex, [3] sampled hemisphere, [4] presence of MRI lesion, [5] presence of daily seizures, [6] number of anti-seizure medications, [7] one- versus two-stage surgery, and [8] biomarker value. Patient and intercept were considered random effects variables. The binary response variable was SOZ or non-SOZ status of electrodes (1 or 0, respectively).

**: In the analysis with only two-stage surgery patients, transfer entropy was calculated using a spectral amplitude time series that was normalized via z-scoring based on the mean and standard deviation of the slow wave sleep period.

***: Bonferroni correction was applied due to the 40 analyses conducted in the main analysis, and significant time points are bolded (Bonferroni corrected p-value < 0.05).

****: The odds ratio was not shown in SPSS due to the enormous effect size.

DF: degrees of freedom; EC = effective connectivity; extraoperative iEEG = extraoperative intracranial electroencephalography; HFO: high-frequency oscillation; intraoperative ECoG = intraoperative electrocorticography; OR: odds ratio; SOZ: seizure onset zone.

**Supplementary Table 9. Binary logistic mixed model characterization of seizure onset zone based on the delta effective connectivity (3 - 4 Hz) response to anesthesia.**

| **Fixed effect variable*** | **Number of patients included in the analysis **** | **DF** | **t-value** | **Uncorrected p-value** | **Bonferroni-corrected p-value***** | **OR** |
| --- | --- | --- | --- | --- | --- | --- |
| Delta EC during Slow Wave Sleep | 11: Two-stage only | 1,279 | -0.054 | 0.957 | > 0.99 | 0.141 |
| **Delta EC during the 1st minute of Sevoflurane**  **(2 vol%)** | **11: Two-stage only** | **941** | **-3.608** | **0.00032** | **0.0128** | **4.048E-27** |
| **Delta EC during the 2nd minute of Sevoflurane**  **(2 vol%)** | **11: Two-stage only** | **928** | **-3.450** | **0.00059** | **0.0236** | **1.727E-27** |
| **Delta EC during the 3rd minute of Sevoflurane**  **(2 vol%)** | **11: Two-stage only** | **643** | **-4.584** | **5.48E-6** | **0.00022** | **3.264E-58** |
| **Delta EC during the 1st minute of Sevoflurane**  **(3 vol%)** | **11: Two-stage only** | **898** | **-5.257** | **1.83E-7** | **7.32E-6** | **1.709E-41** |
| **Delta EC during the 2nd minute of Sevoflurane**  **(3 vol%)** | **11: Two-stage only** | **816** | **-4.799** | **1.90E-6** | **7.6E-5** | **2.743E-33** |
| **Delta EC during the 3rd minute of Sevoflurane**  **(3 vol%)** | **11: Two-stage only** | **547** | **-4.146** | **3.92E-5** | **0.00157** | **1.192E-39** |
| Delta EC during the 1st minute of Sevoflurane  (4 vol%) | 11: Two-stage only | 604 | -2.674 | 0.008 | 0.32 | 5.963E-21 |
| Delta EC during the 2nd minute of Sevoflurane  (4 vol%) | 11: Two-stage only | 538 | -2.124 | 0.034 | > 0.99 | 5.505E-25 |
| Delta EC during the 3rd minute of Sevoflurane  (4 vol%) | 11: Two-stage only | 460 | -0.269 | 0.788 | > 0.99 | 2.816E-4 |

*: The delta effective connectivity value (rated by transfer entropy), as an ECoG/iEEG biomarker, was incorporated into the multivariate binary logistic mixed model analysis, which included eight fixed effect predictors: [1] age, [2] biological sex, [3] sampled hemisphere, [4] presence of MRI lesion, [5] presence of daily seizures, [6] number of anti-seizure medications, [7] one- versus two-stage surgery, and [8] biomarker value. Patient and intercept were considered random effects variables. The binary response variable was SOZ or non-SOZ status of electrodes (1 or 0, respectively).

**: In the analysis with only two-stage surgery patients, transfer entropy was calculated using a spectral amplitude time series that was normalized via z-scoring based on the mean and standard deviation of the slow wave sleep period.

***: Bonferroni correction was applied due to the 40 analyses conducted in the main analysis, and significant time points are bolded (Bonferroni corrected p-value < 0.05).

DF: degrees of freedom; EC = effective connectivity; extraoperative iEEG = extraoperative intracranial electroencephalography; intraoperative ECoG = intraoperative electrocorticography; OR: odds ratio; SOZ: seizure onset zone.

**Supplementary Table 10. Binary logistic mixed model characterization of seizure outcome based on the delta (3-4 Hz) and high-frequency oscillation (80 – 300 Hz) phase-amplitude coupling subtraction value response to anesthesia.**

| **Fixed effect variable*** | **Number of patients included in the analysis** | **DF** | **t-value** | **Uncorrected p-value** | **Bonferroni-corrected p-value**** | **OR** |
| --- | --- | --- | --- | --- | --- | --- |
| Delta-HFO subPAC during Isoflurane  (1 vol%) | 22: One-stage  and Two-stage | 19 | -0.763 | 0.455 | > 0.99 | 0.051 |
| Delta-HFO subPAC during Slow Wave Sleep | 15: Two-stage only | 13 | -1.133 | 0.278 | > 0.99 | 0.003 |
| Delta-HFO subPAC during the 1st minute of Sevoflurane  (2 vol%) | 22: One-stage  and Two-stage | 17 | -0.907 | 0.377 | > 0.99 | 0.099 |
|  | 15: Two-stage only | 12 | -0.474 | 0.644 | > 0.99 | 0.048 |
| Delta-HFO subPAC during the 2nd minute of Sevoflurane  (2 vol%) | 22: One-stage  and Two-stage | 18 | -1.045 | 0.310 | > 0.99 | 0.200 |
|  | 15: Two-stage only | 13 | -0.966 | 0.352 | > 0.99 | 0.241 |
| Delta-HFO subPAC during the 3rd minute of Sevoflurane  (2 vol%) | 22: One-stage  and Two-stage | 14 | -1.026 | 0.322 | > 0.99 | 0.253 |
|  | 15: Two-stage only | 10 | -0.899 | 0.390 | > 0.99 | 0.317 |
| Delta-HFO subPAC during the 1st minute of Sevoflurane  (3 vol%) | 22: One-stage  and Two-stage | 17 | -0.932 | 0.365 | > 0.99 | 0.177 |
|  | 15: Two-stage only | 13 | -0.909 | 0.380 | > 0.99 | 0.215 |
| Delta-HFO subPAC during the 2nd minute of Sevoflurane  (3 vol%) | 22: One-stage  and Two-stage | 17 | -0.878 | 0.392 | > 0.99 | 0.189 |
|  | 15: Two-stage only | 12 | -0.862 | 0.405 | > 0.99 | 0.242 |
| Delta-HFO subPAC during the 3rd minute of Sevoflurane  (3 vol%) | 22: One-stage  and Two-stage | 9 | 0.664 | 0.523 | > 0.99 | 1,685.940 |
|  | 15: Two-stage only | 7 | 0.480 | 0.646 | > 0.99 | 150.448 |
| Delta-HFO subPAC during the 1st minute of Sevoflurane  (4 vol%) | 22: One-stage  and Two-stage | 14 | -0.805 | 0.434 | > 0.99 | 0.259 |
|  | 15: Two-stage only | 10 | -0.751 | 0.470 | > 0.99 | 0.298 |
| Delta-HFO subPAC during the 2nd minute of Sevoflurane  (4 vol%) | 22: One-stage  and Two-stage | 11 | -0.608 | 0.555 | > 0.99 | 0.178 |
|  | 15: Two-stage only | 9 | -0.698 | 0.503 | > 0.99 | 0.249 |
| Delta-HFO subPAC during the 3rd minute of Sevoflurane  (4 vol%) | 22: One-stage  and Two-stage | 8 | -0.813 | 0.440 | > 0.99 | 0.355 |
|  | 15: Two-stage only | 8 | -0.814 | 0.439 | > 0.99 | 0.353 |

*: The subtraction delta-HFO phase-amplitude coupling value (rated by modulation index), as an ECoG/iEEG biomarker, was incorporated into the multivariate binary logistic mixed model analysis, which included eight fixed effect predictors: [1] age, [2] biological sex, [3] sampled hemisphere, [4] presence of MRI lesion, [5] presence of daily seizures, [6] number of anti-seizure medications, [7] one- versus two-stage surgery, and [8] biomarker value. Patient and intercept were considered random effects variables. The binary response variable was ILAE class I seizure freedom or not (1 or 0, respectively). The subtraction biomarker value for each patient was defined as the average of all resected sites minus the average of all retained sites.

**: Bonferroni correction was applied due to the 40 analyses conducted in the main analysis, and significant time points are bolded (Bonferroni corrected p-value < 0.05).

DF: degrees of freedom; extraoperative iEEG = extraoperative intracranial electroencephalography; HFO = high-frequency oscillation; intraoperative ECoG = intraoperative electrocorticography; OR: odds ratio; subPAC: subtraction phase-amplitude coupling.

**Supplementary Table 11. Binary logistic mixed model prediction of seizure outcome based on the high-frequency oscillation effective connectivity subtraction value (80 – 300 Hz) response to anesthesia.**

| **Fixed effect variable*** | **Number of patients included in the analysis **** | **DF** | **t-value** | **Uncorrected p-value** | **Bonferroni-corrected p-value***** | **OR** |
| --- | --- | --- | --- | --- | --- | --- |
| HFO subEC during Isoflurane  (1 vol%) | 22: One-stage  and Two-stage | 19 | -0.530 | 0.602 | > 0.99 | 1.920E-122 |
| HFO subEC during Slow Wave Sleep | 15: Two-stage only | 13 | -1.559 | 0.143 | > 0.99 | **** |
| HFO subEC during the 1st minute of Sevoflurane  (2 vol%) | 22: One-stage  and Two-stage | 16 | -0.349 | 0.732 | > 0.99 | 4.168E-232 |
|  | 15: Two-stage only | 12 | -1.354 | 0.201 | > 0.99 | **** |
| HFO subEC during the 2nd minute of Sevoflurane  (2 vol%) | 22: One-stage  and Two-stage | 17 | 0.131 | 0.897 | > 0.99 | 6.779E55 |
|  | 15: Two-stage only | 13 | -1.326 | 0.208 | > 0.99 | **** |
| HFO subEC during the 3rd minute of Sevoflurane  (2 vol%) | 22: One-stage  and Two-stage | 14 | -1.027 | 0.322 | > 0.99 | **** |
|  | 15: Two-stage only | 10 | -1.229 | 0.247 | > 0.99 | **** |
| HFO subEC during the 1st minute of Sevoflurane  (3 vol%) | 22: One-stage  and Two-stage | 15 | -0.253 | 0.804 | > 0.99 | 1.620E-64 |
|  | 15: Two-stage only | 13 | -1.478 | 0.163 | > 0.99 | **** |
| HFO subEC during the 2nd minute of Sevoflurane  (3 vol%) | 22: One-stage  and Two-stage | 15 | 0.088 | 0.931 | > 0.99 | 2.156E21 |
|  | 15: Two-stage only | 12 | -1.058 | 0.311 | > 0.99 | 1.367E-292 |
| HFO subEC during the 3rd minute of Sevoflurane  (3 vol%) | 22: One-stage  and Two-stage | 9 | 0.623 | 0.549 | > 0.99 | **** |
|  | 15: Two-stage only | 7 | 0.357 | 0.732 | > 0.99 | 1.534E168 |
| HFO subEC during the 1st minute of Sevoflurane  (4 vol%) | 22: One-stage  and Two-stage | 12 | -0.782 | 0.449 | > 0.99 | 1.004E-187 |
|  | 15: Two-stage only | 10 | -1.049 | 0.319 | > 0.99 | 3.536E-257 |
| HFO subEC during the 2nd minute of Sevoflurane  (4 vol%) | 22: One-stage  and Two-stage | 9 | -0.495 | 0.633 | > 0.99 | 1.357E-169 |
|  | 15: Two-stage only | 9 | -0.833 | 0.426 | > 0.99 | 7.136E-129 |
| HFO subEC during the 3rd minute of Sevoflurane  (4 vol%) | 22: One-stage  and Two-stage | 7 | -0.436 | 0.676 | > 0.99 | 7.186E-160 |
|  | 15: Two-stage only | 8 | -0.645 | 0.537 | > 0.99 | 6.795E-110 |

*: The HFO subtraction effective connectivity value (rated by transfer entropy), as an ECoG/iEEG biomarker, was incorporated into the multivariate binary logistic mixed model analysis, which included eight fixed effect predictors: [1] age, [2] biological sex, [3] sampled hemisphere, [4] presence of MRI lesion, [5] presence of daily seizures, [6] number of anti-seizure medications, [7] one- versus two-stage surgery, and [8] biomarker value. Patient and intercept were considered random effects variables. The binary response variable was ILAE class I seizure freedom or not (1 or 0, respectively). The subtraction biomarker value for each patient was defined as the average of all resected sites minus the average of all retained sites.

**: In the analysis including both one-stage and two-stage surgery patients, transfer entropy was calculated using a spectral amplitude time series that was normalized via z-scoring based on the mean and standard deviation of the isoflurane (1 vol%) period. In the analysis with only two-stage surgery patients, transfer entropy was calculated using a spectral amplitude time series that was normalized via z-scoring based on the mean and standard deviation of the slow wave sleep period.

***: Bonferroni correction was applied due to the 40 analyses conducted in the main analysis, and significant time points are bolded (Bonferroni corrected p-value < 0.05).

****: The odds ratio was not shown in SPSS due to the enormous effect size.

DF: degrees of freedom; extraoperative iEEG = extraoperative intracranial electroencephalography; HFO = high-frequency oscillation; intraoperative ECoG = intraoperative electrocorticography; OR: odds ratio; subEC: subtraction effective connectivity.

**Supplementary Table 12. Binary logistic mixed model prediction of seizure outcome based on the delta effective connectivity subtraction value (3 - 4 Hz) response to anesthesia.**

| **Fixed effect variable*** | **Number of patients included in the analysis **** | **DF** | **t-value** | **Uncorrected p-value** | **Bonferroni-corrected p-value***** | **OR** |
| --- | --- | --- | --- | --- | --- | --- |
| Delta subEC during Isoflurane  (1 vol%) | 22: One-stage  and Two-stage | 19 | -0.330 | 0.745 | > 0.99 | 5.773E-107 |
| Delta subEC during Slow Wave Sleep | 15: Two-stage only | 13 | -0.838 | 0.417 | > 0.99 | 9.075E-163 |
| Delta subEC during the 1st minute of Sevoflurane  (2 vol%) | 22: One-stage  and Two-stage | 16 | -0.301 | 0.767 | > 0.99 | 1.143E-46 |
|  | 15: Two-stage only | 12 | -0.753 | 0.466 | > 0.99 | 2.373E-81 |
| Delta subEC during the 2nd minute of Sevoflurane  (2 vol%) | 22: One-stage  and Two-stage | 17 | 0.013 | 0.990 | > 0.99 | 146.946 |
|  | 15: Two-stage only | 13 | -0.358 | 0.726 | > 0.99 | 5.583E-40 |
| Delta subEC during the 3rd minute of Sevoflurane  (2 vol%) | 22: One-stage  and Two-stage | 14 | -0.306 | 0.764 | > 0.99 | 1.306E-33 |
|  | 15: Two-stage only | 10 | -0.737 | 0.478 | > 0.99 | 3.856E-126 |
| Delta subEC during the 1st minute of Sevoflurane  (3 vol%) | 22: One-stage  and Two-stage | 15 | 0.538 | 0.598 | > 0.99 | 2.007E48 |
|  | 15: Two-stage only | 13 | -0.846 | 0.413 | > 0.99 | 1.170E-91 |
| Delta subEC during the 2nd minute of Sevoflurane  (3 vol%) | 22: One-stage  and Two-stage | 15 | 0.356 | 0.727 | > 0.99 | 2.649E50 |
|  | 15: Two-stage only | 12 | -1.176 | 0.262 | > 0.99 | 1.247E-192 |
| Delta subEC during the 3rd minute of Sevoflurane  (3 vol%) | 22: One-stage  and Two-stage | 9 | 0.241 | 0.815 | > 0.99 | 1.473E60 |
|  | 15: Two-stage only | 7 | -0.884 | 0.406 | > 0.99 | **** |
| Delta subEC during the 1st minute of Sevoflurane  (4 vol%) | 22: One-stage  and Two-stage | 12 | 0.810 | 0.434 | > 0.99 | 2.375E100 |
|  | 15: Two-stage only | 10 | -0.613 | 0.554 | > 0.99 | 7.388E-56 |
| Delta subEC during the 2nd minute of Sevoflurane  (4 vol%) | 22: One-stage  and Two-stage | 9 | 0.111 | 0.914 | > 0.99 | 5.723E10 |
|  | 15: Two-stage only | 9 | -0.711 | 0.495 | > 0.99 | 2.565E-77 |
| Delta subEC during the 3rd minute of Sevoflurane  (4 vol%) | 22: One-stage  and Two-stage | 7 | 1.264 | 0.247 | > 0.99 | 1.519E208 |
|  | 15: Two-stage only | 8 | -0.259 | 0.802 | > 0.99 | 4.798E-25 |

*: The delta subtraction effective connectivity value (rated by transfer entropy), as an ECoG/iEEG biomarker, was incorporated into the multivariate binary logistic mixed model analysis, which included eight fixed effect predictors: [1] age, [2] biological sex, [3] sampled hemisphere, [4] presence of MRI lesion, [5] presence of daily seizures, [6] number of anti-seizure medications, [7] one- versus two-stage surgery, and [8] biomarker value. Patient and intercept were considered random effects variables. The binary response variable was ILAE class I seizure freedom or not (1 or 0, respectively). The subtraction biomarker value for each patient was defined as the average of all resected sites minus the average of all retained sites.

**: In the analysis including both one-stage and two-stage surgery patients, transfer entropy was calculated using a spectral amplitude time series that was normalized via z-scoring based on the mean and standard deviation of the isoflurane (1 vol%) period. In the analysis with only two-stage surgery patients, transfer entropy was calculated using a spectral amplitude time series that was normalized via z-scoring based on the mean and standard deviation of the slow wave sleep period.

***: Bonferroni correction was applied due to the 40 analyses conducted in the main analysis, and significant time points are bolded (Bonferroni corrected p-value < 0.05).

****: The odds ratio was not shown in SPSS due to the enormous effect size.

DF: degrees of freedom; extraoperative iEEG = extraoperative intracranial electroencephalography; intraoperative ECoG = intraoperative electrocorticography; OR: odds ratio; subEC: subtraction effective connectivity.
